# ProteomeLM: A proteome-scale language model enables accurate and rapid prediction of protein-protein interactions and gene essentiality across taxa

**DOI:** 10.1101/2025.08.01.668221

**Authors:** Cyril Malbranke, Gionata Paolo Zalaffi, Anne-Florence Bitbol

## Abstract

Language models trained on biological sequences are advancing inference tasks from the scale of single proteins to that of genomic neighborhoods. Here, we introduce ProteomeLM, a transformer-based language model that uniquely operates on entire proteomes from species spanning the tree of life. ProteomeLM is trained to reconstruct masked protein embeddings using the whole proteomic context, yielding contextualized protein representations that reflect proteome-scale functional constraints. Notably, ProteomeLM’s attention coefficients encode protein-protein interactions (PPI), despite being trained without interaction labels. Furthermore, it enables interactome-wide PPI screening that is substantially more accurate, and orders of magnitude faster, than amino-acid coevolution-based methods. We further develop ProteomeLM-PPI, a supervised model that combines ProteomeLM embeddings and attention coefficients to achieve state-of-the-art PPI prediction across benchmarks and species. Finally, we introduce ProteomeLM-Ess, a supervised gene essentiality predictor that generalizes across diverse taxa. Our results demonstrate the potential of proteome-scale language models for addressing function and interactions at the organism level.

**Significance statement:** Predicting protein interactions and functions is a key challenge in biology. Although deep learning-based language models are advancing the analysis of individual protein sequences and of genomic neighborhoods, they struggle to capture properties involving all the proteins expressed in a cell, such as protein–protein interactions (PPI) and gene essentiality. We present ProteomeLM, a language model that reasons on entire proteomes across diverse species. ProteomeLM captures PPI without supervision, and enables more accurate and faster screening of entire interactomes than current sequence-based approaches. ProteomeLM also delivers state-of-the-art supervised PPI prediction, and improves supervised prediction of gene essentiality compared to protein language models. These results demonstrate the potential of proteome-scale language models to reveal system-level organization and functional relationships.

## Introduction

Recently, deep learning approaches have brought important progress to inference from biological sequence data. Protein language models are deep learning models based on natural language processing methods. Trained on large ensembles of protein sequences, they learn sequence representations that encode structural and functional signals [1–12], and have advanced the prediction of protein structure [9, 10], subcellular localization [13], and mutational effects [14, 15]. Similarly, genome language models [16–26] have given insight in non-coding DNA, gene expression and taxonomic classification [17–19], capturing operons, enzymatic function [21] and within-operon protein-protein interactions (PPI) [22], and predicting mutation effects [20]. However, so far, these models span at most hundreds of kilobases or a few megabases [18, 19, 23–26]. As they do not capture dependencies across entire genomes, especially in eukaryotes, these models cannot predict system-level properties such as PPI involving distant genomic regions.

PPI are fundamental to most biological processes, including signal transduction, cellular metabolism, and immune responses. Knowing these interactions is critical for deciphering cellular processes and for developing therapeutic interventions. However, large-scale PPI determination – up to the full PPI networks of non-model species – remains a significant challenge [27]. Indeed, precise experimental methods are both labor-intensive and costly, particularly when scaled to entire proteomes, and high-throughput ones have limited accuracy [28]. While curated PPI databases have grown [29–31], they remain incomplete and biased toward well-studied species and interaction types. Computational PPI predictions have been developed to overcome this gap. Structure-based ones, including docking [32–35] and multimeric folding algorithms like AlphaFold-Multimer [36], have achieved remarkable accuracy for specific interactions [37], but remain computationally intensive. Sequence-based methods, which rely on evolutionary signals, offer better scalability. Amino acids that are in contact at the interface of protein complexes need to maintain physico-chemical complementarity through evolution, yielding correlations in amino-acid usage at contacting sites, known as coevolution [38–41]. This signal can be detected by Potts models, also known as direct coupling analysis (DCA), which are trained on multiple sequence alignments of homologous proteins for each candidate PPI [42–48]. At a larger scale, during evolution, genes coding for proteins that interact tend to undergo similar evolutionary pressures and to be either present or absent together in each genome, thus having correlated co-occurence patterns across genomes. This coevolution between genes is exploited by phylogenetic profiling [49–57] and co-occurrence methods [58]. While such coevolution methods are often effective in bacteria and other well-represented clades, they struggle in eukaryotes or poorly sampled taxa, and require careful curation of orthologs, and for DCA, paired multiple sequence alignments [45, 46, 59, 60]. Protein language models trained with the masked language modeling (MLM) objective of predicting masked amino acids in sequences using the surrounding context [3, 9–12] learn coevolution between amino acids. This allows them, in particular those based on multiple sequence alignments [9], to capture protein structure, an ability exploited in AlphaFold2 via its EvoFormer module [61]. Given the success of protein language models at capturing coevolution between amino acids, it is tantalizing to develop such models at the proteome scale, i.e. taking as input the ensemble of proteins encoded by a genome. Indeed, we posit that such models should capture coevolution between proteins, thereby generalizing over phylogenetic profiling methods [49–57]. This should make them highly suited to provide predictions of complete protein-protein interaction networks, and of their evolution. Importantly, once trained, the computational cost of prediction on new species should be small. Moreover, such foundation models can also be used for other downstream applications where proteome context information is important, such as predicting gene essentiality [62].

In this paper, we introduce ProteomeLM, a transformer-based language model that uniquely reasons on entire proteomes from multiple species spanning the tree of life. ProteomeLM leverages embeddings from the protein language model ESM-Cambrian [12], and is thus informed of functional protein-level properties, which it integrates at the proteome scale. We show that ProteomeLM’s attention coefficients learn PPI in an unsupervised way. These PPI include broad functional relationships and protein complex membership, which yield distinct signals. Building on this ability of ProteomeLM, we propose a new method to screen whole interactomes orders of magnitude faster than DCA pipelines, and we demonstrate that it substantially outperforms them. Next, we introduce ProteomeLM-PPI, a supervised PPI prediction network that combines ProteomeLM embeddings and attention coefficients, and we show that it achieves state-of-the-art supervised PPI prediction across species and benchmarks. Finally, as another example of downstream tasks allowed by ProteomeLM embeddings, we introduce ProteomeLM-Ess, a supervised predictor of gene essentiality that generalizes across diverse taxa. Our results show that representing proteins in their full proteome context allows ProteomeLM to capture system-level biological signals that remain inaccessible to models that reason on individual proteins or on local genomic contexts.

## Results

### A proteome-scale language model leveraging protein language model representations

We introduce ProteomeLM, a transformer-based Proteome Language Model (LM). We trained ProteomeLM on a large corpus of proteomes, spanning the tree of life, from bacteria and archaea to eukaryotes and viruses, to learn protein representations in the context of complete proteomes. ProteomeLM takes as input a proteome, i.e. the set of proteins encoded by a given genome, and aims to capture the functional and evolutionary signals present between proteins at the proteome level.

Specifically, we start from each protein’s amino-acid sequence, and represent it by an embedding generated by the protein language model ESM-Cambrian (ESM-C) [12], see Figure 1 and “Protein Representations” in Methods. This allows our model to leverage the rich functional sequence-derived properties [63, 64] learned for each protein by protein language models [3, 9, 10, 14] (see also [21, 22, 65]). During training, a subset of these protein embeddings is masked, and the model is tasked with reconstructing them using the remaining unmasked protein embeddings from the same proteome, see Figure 1. This masked language modeling prediction task allows ProteomeLM to learn the dependencies between proteins encoded by a given genome.

**Figure 1:**
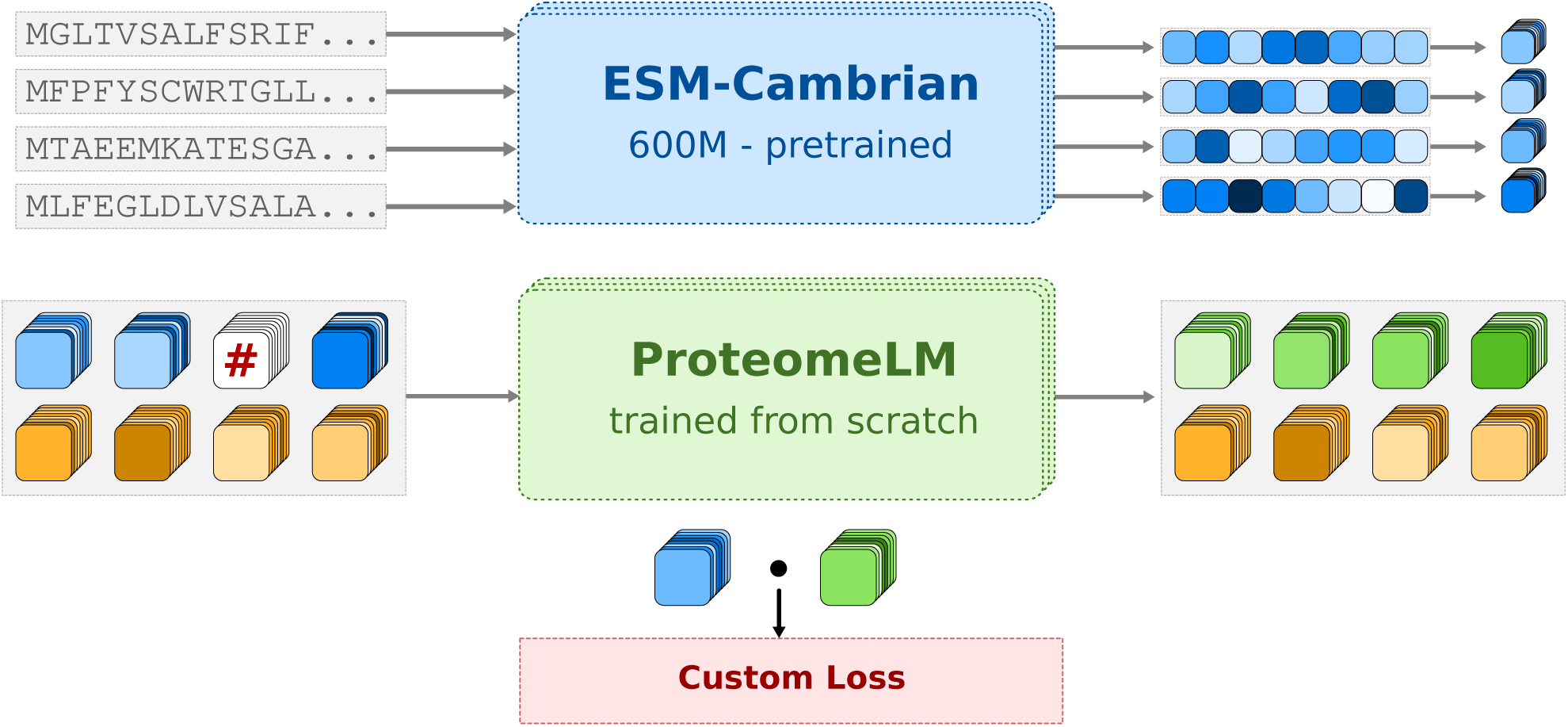
ProteomeLM training. Input amino-acid sequences, corresponding to proteins encoded by a genome, are embedded through the pretrained ESM-C model, yielding a fixed-dimensional embedding for each protein. The embeddings serve as input to ProteomeLM, trained from scratch to predict the masked embeddings of proteins in the context of their proteome (see Methods). Masked embeddings are indicated by “#” in the schematic. Proteins are annotated by a functional encoding (shown in orange below embeddings) representing their orthologous group. ProteomeLM’s training uses a custom polar loss (see Methods): for each masked protein, it essentially aims to minimize the difference between the ESM-C embedding and the ProteomeLM embedding, but in a protein family-specific manner.

A challenge when constructing a model that can reason across diverse organisms is the fact that their genome organization is very different. For instance, while bacterial genes are organized in operons, functionally related proteins are often not encoded in proximity in eukaryotic genomes, and gene order is less conserved in eukaryotes than in prokaryotes. Therefore, ProteomeLM does not employ positional encoding along the genome, which sets it apart from existing genome language models (and protein language models). Instead, we propose a *functional encoding*, based on orthology, which captures shared evolutionary and functional relationships across genes in different genomes (see “Functional Encoding” in Methods). Orthologous groups comprise genes in different species that descend from a common ancestral gene via speciation, and generally have retained the same function. Phylogenetic profiling methods have shown that the presence and absence data of orthologous groups, across genomes, contain information about functional relationships [49–51, 54, 66]. For instance, genes coding for interacting proteins tend to have correlated presence-absence vectors across genomes, allowing to predict interactions. Hence, we posit that functional encoding will help ProteomeLM to learn coevolution between proteins in a proteome, and thus, their functional relationships. In practice, orthologous groups were collected from OrthoDB [67]. Note that they were built using statistical homolog matching, and no human annotations.

ProteomeLM was trained on nearly 32,000 annotated proteomes, spanning all domains of life (see “Dataset” in Methods). By learning to reconstruct masked proteins from their genomic context, ProteomeLM is encouraged to learn the relationships that define protein function and interactions. We trained versions of ProteomeLM with multiple sizes, ranging from 6M to 328M parameters. We observed stable learning dynamics in all cases (see “Training dynamics and scaling behavior” in Methods).

### ProteomeLM attention coefficients capture protein-protein interactions

ProteomeLM was trained on full proteomes, and informed by protein-level properties, thanks to the use of embeddings from protein language models. Does it learn protein-protein interactions (PPI) by being trained to predict a protein’s embedding using the context of its proteome? To address this question, we examine the attention coefficients of the model [5, 9, 68]. Indeed, in a transformer model, attention coefficients encode the importance of each part of the context (here, of each protein) when performing masked language modeling (here, when aiming to predict another masked protein). Thus, we hypothesize that these coefficients should capture functional dependencies between proteins, such as PPI. For each pair of protein in five species, we compare these attention coefficients to known PPI. The species considered are *Escherichia coli*, *Saccharomyces cerevisiae*, *Caenorhabditis elegans*, *Drosophila melanogaster*, *Mus musculus*, and *Homo sapiens*. Specifically, we use the D-SCRIPT dataset [27], which is derived from the STRING database [69], and focuses exclusively on experimentally validated physical interactions. Complete proteomes from the five species considered were processed through ProteomeLM, and we extracted attention coefficients for each positive and negative interaction pair across both datasets.

Figure 2A-C shows that many attention heads of ProteomeLM are predictive of interaction labels. Moreover, we observe that several attention heads possess significant predictive power across all species considered (3 extra species are shown in Figure S6). In particular, head 7 of layer 3 achieves an AUC of 0.92 in *E. coli*, while also performing strongly in other species. The larger ProteomeLM-M also exhibits strong unsupervised PPI recovery, see Figure S7. Thus, ProteomeLM can identify PPI among vast numbers of possible protein pairs in a complete proteome (e.g., ∼ 4,000 proteins in *E. coli* and ∼ 20,000 in humans, leading to ∼ 8 × 10^6^ and ∼ 2 × 10^8^ possible pairs, respectively) in an unsupervised manner, without any fine-tuning. This is especially compelling given that ProteomeLM does not rely on gene order or local genomic context. The learning of PPI arises directly from the masked prediction training, which promotes the learning of dependencies between proteins in a proteome. Our result is reminiscent of the finding that protein language models’ attention coefficients carry information about residue-residue interactions involved in the three-dimensional structure of proteins, while being trained only on sequence data with the objective of filling in masked amino acids [3, 9, 10].

**Figure 2:**
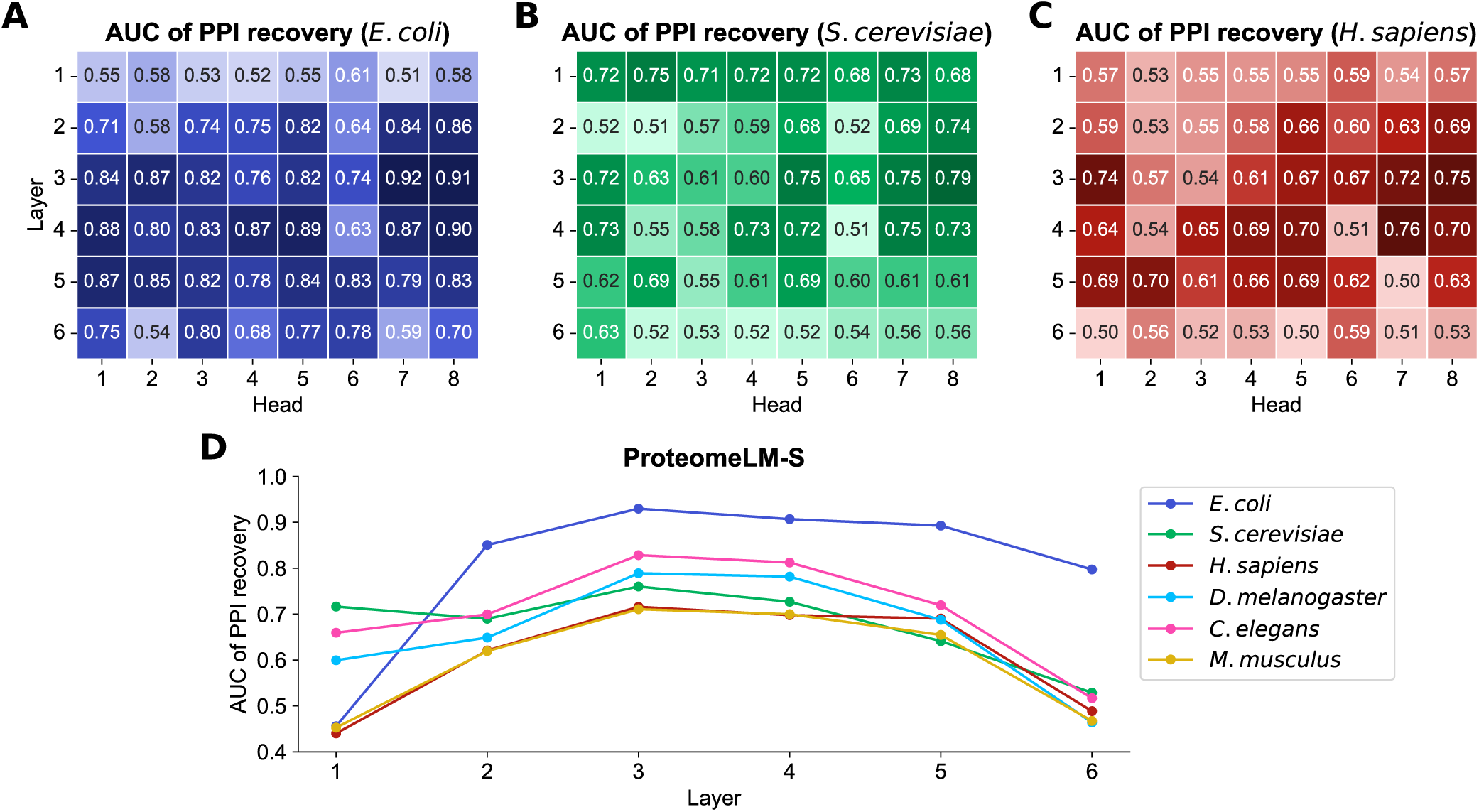
Unsupervised detection of PPI using ProteomeLM attention coefficients. We assess the ability of the attention coefficients of ProteomeLM-S (36M parameters) at predicting PPI from the D-SCRIPT dataset [27], using the Area Under the Receiver Operating Characteristic Curve (AUC) as a metric. **(A-C)** In three species, we report the AUC of each attention head in each layer of ProteomeLM, measuring its ability to distinguish interacting from non-interacting protein pairs (1: perfect classifier; 0.5: random). **(D)**: We show the AUC obtained for the mean of attention coefficients over all heads in each layer of ProteomeLM-S, in each of the six species considered.

Figure 2D further shows that PPI are most accurately captured by attention heads in the central layers of the model. A similar trend is observed across other model sizes (XS, M, and L), see Figure S8. In protein language models, central layers are known to capture more complex interactions than early layers [5]. Thus, our finding supports the notion that higher-order interactions, rather than simple local features, are essential for understanding PPI, and that ProteomeLM can extract such complex biological signals.

We next ask what types of PPI are captured by ProteomeLM’s attention heads. As ProteomeLM learns statistical dependencies between proteins from their context across thousands of proteomes, and shares similarities with phylogenetic profiling [49], its attention coefficients are expected to capture broader functional associations (e.g., co-regulation) beyond direct physical binding. To address this question, we build a benchmark distinguishing direct interactions (proteins that directly bind to each other, from PDB [70]), same-complex interactions (protein pairs found in the same complex but not directly binding, from PDB), and genetic associations (co-expression pairs from STRING [69]). We find that ProteomeLM captures all three interaction types in its attention coefficients across *E. coli*, *S. cerevisiae* and *H. sapiens*. Genetic associations are recovered with higher accuracy (AUC ≥ 0.92 in all three species) than direct or same-complex interactions (0.74 ≤ AUC ≤ 0.92, see Supplementary Section 1). Moreover, a linear classifier based on its attention coefficients can distinguish direct interactions from genetic associations with high accuracy (AUC = 0.89 in yeast, 0.90 in humans), although the signal separating direct from same-complex interactions is less strong (see Supplementary Section 1). Thus, ProteomeLM learns representations that disentangle physical binding from functional associations. As a specific test of ProteomeLM’s ability to distinguish direct interactions from same-complex interactions, we further consider two structurally resolved protein complexes: the *E. coli* ribosome and the *S. cerevisiae* TRiC/CCT chaperonin. In both cases, ProteomeLM excels at distinguishing protein pairs belonging to a given complex from the rest of the proteome (AUC ≥ 0.99). Direct interactions within the complexes are more weakly captured, with statistically significant signal for the *E. coli* ribosome but not for the *S. cerevisiae* TRiC/CCT chaperonin (see Supplementary Section 2). The latter complex is particularly challenging, as all of its subunits descend from a common ancestor.

Overall, ProteomeLM is an excellent predictor of broad functional relationships and of protein complex membership. Furthermore, it outperforms its input ESM-C embeddings and functional encodings, which already encode evolutionary and functional information (see Supplementary Section 3).

### ProteomeLM provides fast and accurate PPI screening

ProteomeLM successfully captures PPI, and was trained on organisms spanning the tree of life. Can it thus contribute to shedding light on complete interactomes in various species? Current large-scale interactome prediction workflows generally rely on a two-stage pipeline [48, 71, 72]. First, relatively light sequence-based methods exploiting coevolution in multiple sequence alignments (MSAs), in particular Potts models, also known as Direct Coupling Analysis (DCA) [42–44], are used to identify promising protein pairs that may interact. Second, heavier structure-based methods like AlphaFold-Multimer [36] or RoseTTAFold-PPI [72] are used to further analyze these candidate pairs, through computationally intensive structural modeling. While effective, the first step of this approach is limited by the computational cost of MSA generation, and by the sheer number of pairwise models required to scan large proteomes. Indeed, DCA is a family-specific model, so one model has to be trained per candidate protein pair. To this day, such large-scale PPI screens have thus only been performed on a few model organisms, namely *E. coli* [47] and *S. cerevisiae* [48], and very recently on *H. sapiens* [72], and on 19 human pathogens [71].

To assess ProteomeLM’s promise as a first filter for interactome prediction, we trained a lightly-supervised and lightweight classifier on attention coefficients from ProteomeLM, and evaluated its performance on the full human interactome and across pathogen interactomes. We compared its computational requirements and its performance to those reported in a recent large-scale study that applied DCA respectively to over 190 million *Homo sapiens* protein pairs [72]. We also tested our method on 19 human bacterial pathogens considered in [71], collectively comprising 102 million protein pairs. Specifically, our classifier is a logistic regression aiming to predict PPI from a combination of the 48 attention heads of ProteomeLM-S (see Figure 2A-C), and is trained over a small set of 100 interacting pairs and 1,000 random pairs (treated as non-interacting), which are later held out from evaluation. Positive pairs are sampled among the pairs that have a confidence score above 0.99 (extremely high) in the STRING database [69], either among human pairs or among pathogenic pairs in the corresponding datasets.

In Ref. [72], the DCA pipeline required over 30 days on 50–100 GPUs to process the human proteome. In contrast, ProteomeLM inference, including ESM-C embeddings and attention coefficient computation, takes under 10 minutes per proteome (e.g., *H. sapiens*) on a single RTX A6000 GPU. Importantly, these features are calculated for all possible protein pairs, without the need to train a separate model for each candidate pair. Moreover, ProteomeLM training was completed in 3 days on a single H100 GPU, and generalizes to all downstream tasks without retraining. As shown in Figure 3A, ProteomeLM thus reduces overall compute time by up to six orders of magnitude for inference alone, and by three orders of magnitude when training is included (see Supplementary Section 4 for details).

**Figure 3:**
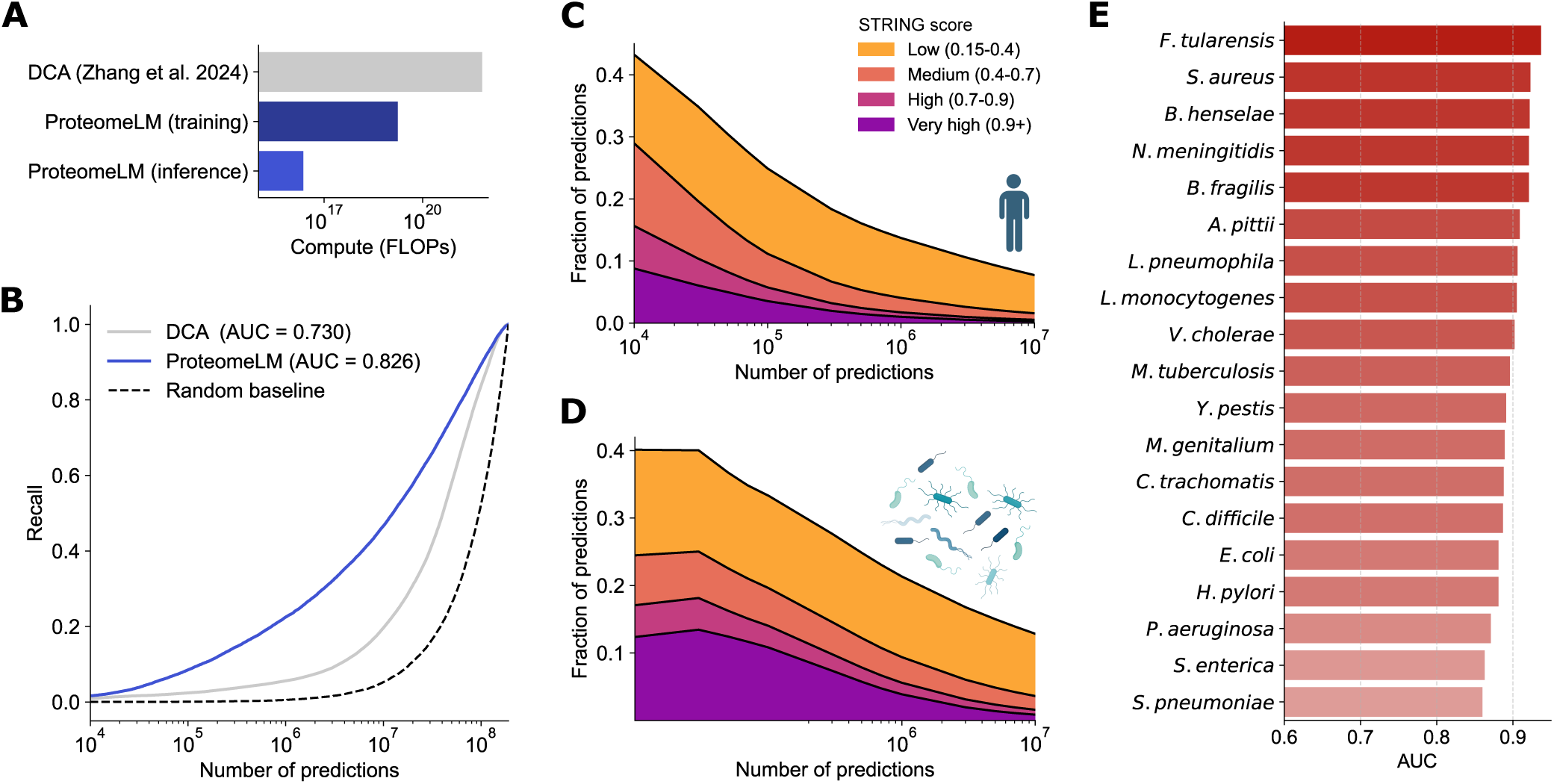
Fast and high-precision screening of whole interactomes using ProteomeLM. **(A)** Compute required to analyze the full human proteome using ProteomeLM versus DCA. We compare the compute needed to train 190 million DCA models in order to screen the *H. sapiens* proteome [72] with the total compute we used to train ProteomeLM (which was done once across a large number of organisms) and with the compute required to apply ProteomeLM to the *H. sapiens* proteome (inference). ProteomeLM offers a speed-up of up to 6 orders of magnitude when focusing on inference, and of 3 orders of magnitude when including training. **(B)** *H. sapiens* interactome recovery by ProteomeLM and by DCA. Recall is shown versus the number of predictions made. Predictions are made by ranking all possible protein pairs by their score, either from Proteome or from DCA [72]. **(C–D)** Fraction of top-scoring predictions that correspond to known interactions in the STRING database, for *H. sapiens* (C) and 19 human bacterial pathogens (D). **(E)** Interactome recovery performance across 19 pathogenic bacterial species, measured by the area under the receiver operating characteristic curve (AUC).

Moreover, Figure 3B shows that ProteomeLM significantly outperforms DCA in recovering experimen-tally validated interactions. In *H. sapiens*, ProteomeLM achieves an area under the receiver operating characteristic curve (AUC) of 0.83, compared to 0.73 for DCA. Among the top 10 million scored pairs, it recovers 50% of known PPI, versus only 20% for DCA. At higher precision thresholds (e.g., top 1 million), ProteomeLM also delivers improved recall. Thus, ProteomeLM has a very strong potential to reduce the burden on downstream structure-based modeling for precise PPI prediction.

We further examine the overlap between ProteomeLM’s top predictions and STRING [69] annotations. Figure 3C shows that in *H. sapiens*, over 40% of the top 10,000 predictions align with known or suspected interactions, and nearly 10% correspond to high-confidence interactions according to STRING. This suggests that ProteomeLM captures meaningful biological associations, even beyond the most confidently labeled pairs.

We extend our analysis to 19 human bacterial pathogens [71]. Figure 3D shows that more than 40% of the top 10,000 predictions are supported by STRING. The per-species breakdown is shown in Supplementary Figure S9, revealing consistent patterns of STRING support of ProteomeLM predictions across diverse pathogens. Figure 3E further reports AUC values for each of the 19 species, ranging from 0.87 to 0.92. These results confirm that ProteomeLM generalizes well and maintains consistently strong performance across highly diverse taxa.

Together, these findings demonstrate that ProteomeLM is a fast, scalable, and accurate framework for interactome screening. Its ability to recover known interactions and predict plausible novel ones, combined with strongly reduced computational costs, demonstrates its strong potential as a replacement for amino-acid coevolution-based filters in proteome-scale PPI prediction pipelines. ProteomeLM therefore opens the way to high-throughput interactome inference across species, including for organisms lacking curated interaction data.

### ProteomeLM delivers state-of-the-art supervised PPI prediction

Since ProteomeLM learns PPI through its attention coefficients without using interaction labels (see Figure 2), and can provide accurate and fast PPI screening, it is tantalizing to further exploit this signal. Can features from ProteomeLM advance supervised sequence-based PPI prediction? To address this question, we introduce ProteomeLM-PPI, a new supervised PPI prediction network that employs both node-type features (i.e. embeddings from ESM-C and ProteomeLM for individual proteins) and edge-type features (i.e. ProteomeLM attention values for pairs of proteins), see Figure 4A.

**Figure 4:**
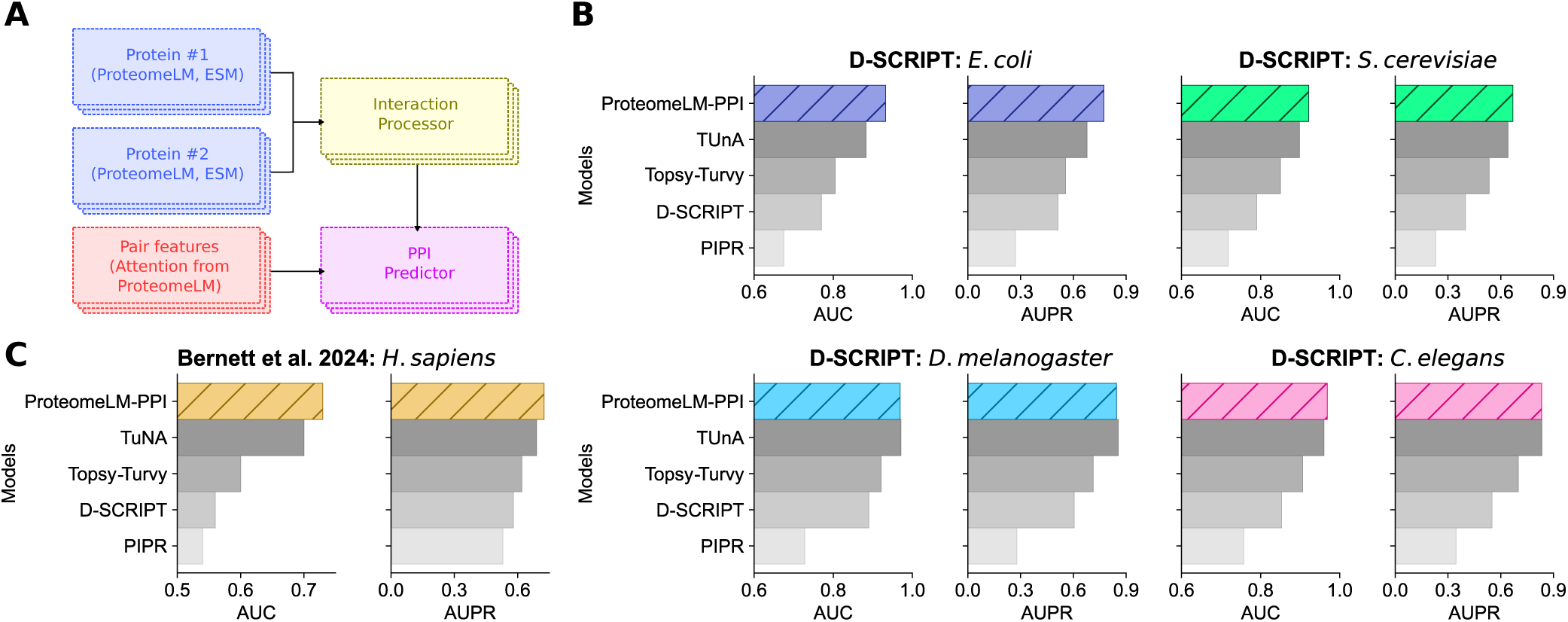
Supervised prediction of PPI using ProteomeLM. **(A)** Architecture of the supervised model trained to predict PPI. The model comprises four components. For each protein in a candidate pair, its representations from ProteomeLM-S and ESM-C are independently processed through a feature encoder (blue), which is a feedforward neural network with two hidden layers. The resulting node features are combined in an interaction processor (yellow) that captures joint information. In parallel, edge features from ProteomeLM attention coefficients are processed through a separate module (red). The final PPI predictor (pink) is a classifier that integrates both node- and edge-level features to predict the interaction probability. **(B)** Cross-species generalization performance on the D-SCRIPT dataset [27], compared to state-of-the-art methods [27, 73–75]. All models are trained and validated on human PPI and tested on other species, and results are averaged over five technical replicates. **(C)** Performance on the dataset from [76], averaged over five technical replicates and compared to state-of-the-art methods. In panels B and C, results are provided in terms of area under the receiver operating characteristic curve (AUC) and area under the precision-recall curve (AUPR).

We trained and evaluated ProteomeLM-PPI on two PPI datasets. The first one is the multi-species D-SCRIPT dataset [27], already used in the previous sections. The second one is a human-specific dataset which was designed to address common biases in the training, validation, and test splits of previous PPI benchmarks, in particular leakage [76]. For both datasets, we used the same training, validation, and test splits as in the recent TUnA method [75]. ProteomeLM-PPI was trained on the training set, with early stopping based on validation performance to avoid overfitting.

In the D-SCRIPT dataset, the training set is composed of human PPI and the validation set comprises held-out human PPI [75]. Testing on PPI from other species thus allows to assess cross-species generalization power. Figure 4B shows that ProteomeLM-PPI outperforms state-of-the-art methods on *E. coli* and *S. cerevisiae*, and performs comparably to them on *D. melanogaster* and *C. elegans*. In particular, it leads to an AUPR improvement of more than 0.1 (from 0.67 to 0.79) over the previous state of the art, TUnA [75], on *E. coli*. These results highlight ProteomeLM’s strong ability to capture PPI signals, its robustness across diverse branches of the tree of life, and its ability to generalize from one species to other ones.

In Figure 4C, we report results on the dataset from Ref. [76]. We observe that ProteomeLM-PPI yields very strong AUC and AUPR scores, consistently reaching or outperforming state-of-the-art methods including TUnA, which employed ESM-2 embeddings. This result shows the robustness of ProteomeLM-PPI to biases of PPI prediction benchmarks [76].

To better understand the contribution of each feature used by ProteomeLM-PPI, we evaluate different combinations of these features in Supplementary Figure S10. We observe that both ProteomeLM embeddings and attention coefficients are individually informative, and that their combination consistently yields the best predictive performance. This suggests that embeddings and attention coefficients contain complementary information that is synergistic for PPI prediction.

### ProteomeLM improves gene essentiality prediction

While we focused on PPI prediction so far, ProteomeLM is a foundation model that can be used for diverse tasks where proteome-level information is important. Here, we consider another important task, which consists in predicting which genes are essential, i.e. necessary for survival or reproduction of an organism. Both protein or gene sequence on the one hand, and genomic context and protein-protein interactions on the other hand, have been found to matter for predicting essentiality [62]. Can ProteomeLM, which exploits these diverse elements, advance gene essentiality prediction?

To address this question, we introduce ProteomeLM-Ess, a supervised predictor of gene essentiality that takes as input ProteomeLM embeddings. To train and test this classification model, we used the OGEE database [77], which collects gene essentiality data from 127 experimental studies, spanning 91 species, for a total of 213,608 labeled genes. To create the training, validation and test split, we clustered proteins based on sequence similarity (see Methods). ProteomeLM-Ess is a two-layer fully connected network, and can take as input ProteomeLM embeddings from any layer.

Figure 5A shows the performance of ProteomeLM-Ess versus the depth of the layer whose embeddings are used as its input, for all ProteomeLM sizes. This performance is also compared to that of a similar classifier that starts from ESM-C embeddings instead of ProteomeLM ones. We observe that classifiers based on ProteomeLM embeddings significantly outperform those based on ESM-C embeddings. This demonstrates that the contextualized whole proteome-aware information present in ProteomeLM embeddings allows to better capture gene essentiality than protein-level information. Besides, we observe that the embeddings from intermediate layers of ProteomeLM produce the best gene essentiality classifiers, which is consistent with our observations for unsupervised PPI prediction (see Figure 2D). Furthermore, the performance of ProteomeLM-Ess scales with ProteomeLM size. This suggests that larger models are able to encode more information about gene essentiality into their representations. Interestingly, larger sizes appear to be more useful for gene essentiality than for PPI prediction (see Figure S9). Overall, the best performing version of ProteomeLM-Ess is the one trained on the embeddings of layer 8 of ProteomeLM-L, yielding an AUC of 0.93.

**Figure 5:**
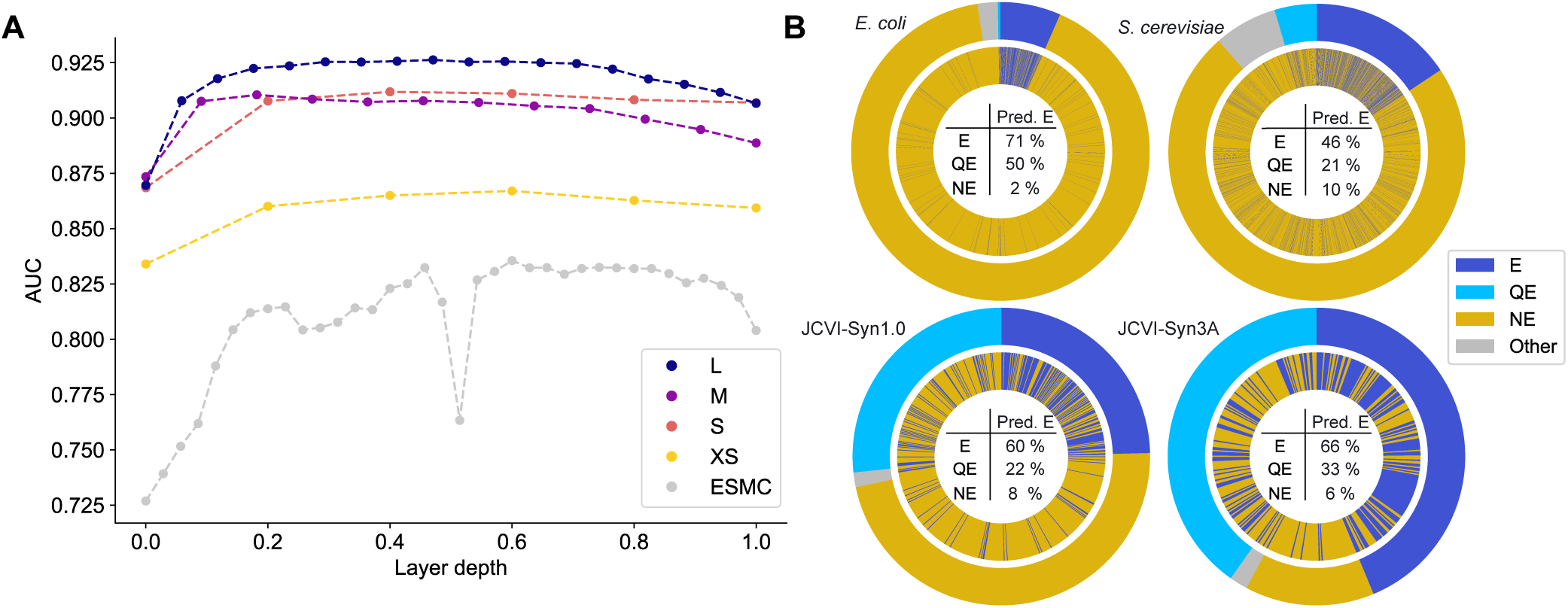
Gene essentiality prediction with ProteomeLM-Ess. **(A)** Area under the ROC curve (AUC) for binary classifiers that take as input embeddings from each layer of ProteomeLM (ProteomeLM-Ess) or ESM-C. To facilitate comparison between models with different sizes, the AUC is plotted versus the normalized layer depth, i.e. the ratio of the layer index to the number of layers. **(B)** Comparison between ProteomeLM-Ess predictions (inner rings) and experimental essentiality labels (outer rings) for *E. coli* K-12, *S. cerevisiae*, and the synthetic cells JCVI-Syn1.0 and JCVI-Syn3A. ProteomeLM-Ess uses embeddings from layer 8 of ProteomeLM-L. These genomes are not in the training set of ProteomeLM-Ess. For each of them, we predict as essential the top *N* genes in terms of ProteomeLM-Ess score, where *N* is the number of genes experimentally labeled essential. While ProteomeLM-Ess classifies genes as essential (E) or non-essential (NE), some genes in these species (many in minimal cells) are labeled as quasi-essential (QE). The percentage of genes labeled as E, QE or NE that are predicted as E by ProteomeLM-Ess is indicated in each case. AUC values: *E. coli*, 0.95; *S. cerevisiae*, 0.81; JCVI-Syn1.0, 0.88; JCVI-Syn3A, 0.83.

Can ProteomeLM-Ess generalize to unseen proteomes? To address this question, we held out the entire proteomes of *S. cerevisiae* and of *E. coli* during training. Figure 5B (top) shows the performance of ProteomeLM-Ess on these two held-out proteomes. We obtain good results, especially in *E. coli*, where 71% (resp. 2%) of genes experimentally determined to be essential (resp. non-essential) were correctly predicted as essential (resp. non-essential) by ProteomeLM-Ess. To further explore generalization ability, we collected essentiality data for the synthetic cells JCVI-Syn1.0 [78] and JCVI-Syn3A [79, 80], which are not present in OGEE. Moreover, JCVI-Syn3A has a genome that is engineered to be close to minimal, so it is both far from genomes in the training set and a stringent test of gene essentiality prediction. Figure 5B (bottom) shows that we obtain good performance in these cases too, even for JCVI-Syn3A. Hence, ProteomeLM-Ess is able to generalize to unseen taxa.

We further compare ProteomeLM-Ess to several existing gene essentiality prediction methods [24, 25, 81–88] on *E. coli* and *S. cerevisiae*, which were both held out from the training set. Overall, ProteomeLM-Ess achieves state of the art performance for *E. coli*. It is also state of the art for *S. cerevisiae* among methods trained without labeled data from that species. Methods trained using labeled data from *S. cerevisiae*, however, perform as well as or better than ProteomeLM-Ess (see Supplementary Section 5).

## Discussion

We introduced ProteomeLM, a transformer-based language model that learns contextualized protein representations from complete proteomes spanning the whole tree of life. Trained to reconstruct masked protein representations from the other proteins of a proteome, ProteomeLM learns dependencies between proteins that reflect functional and evolutionary constraints. This makes it well-suited to predict proteome-scale properties. We showed that ProteomeLM captures PPI in its attention heads, without any supervision. Specifically, ProteomeLM is an excellent predictor of both broad functional relationships and protein complex membership, and can distinguish them.

We then demonstrated that ProteomeLM can be used as a highly scalable and effective tool for whole-interactome screening, yielding substantially higher accuracy than DCA, with a computational cost reduced by up to 6 orders of magnitude. This makes ProteomeLM a strong choice for large-scale PPI screening tasks, including in poorly annotated organisms. Next, we designed ProteomeLM-PPI, a supervised PPI prediction network that leverages both embeddings and attention coefficients from ProteomeLM, and achieves state-of-the-art performance across species and benchmarks. Finally, we trained ProteomeLM-Ess, a supervised classifier for gene essentiality prediction that starts from ProteomeLM embeddings. We showed that classifiers trained on ProteomeLM embeddings outperform those trained on ESM-C embeddings for this task, confirming that ProteomeLM’s contextualized embeddings encode biologically relevant proteome-scale information.

In summary, ProteomeLM is a novel framework that learns whole proteome-aware protein representations across the tree of life. As a foundation language model trained using the MLM objective, ProteomeLM can be used for many downstream tasks. The possibility of using the pre-trained ProteomeLM, and even of employing pre-computed ProteomeLM embeddings, means that using it in inference for downstream tasks can be very computationally efficient. We demonstrated that ProteomeLM reaches very strong performance on tasks ranging from PPI screening to gene essentiality prediction, and combines speed, interpretability, and accuracy. We expect that proteome-aware language models will become increasingly important in the coming years, enabling new ways of modeling system-level biological properties at the scale of proteomes and cells.

Integrating local genomic context to protein language model embeddings has proved very useful for function inference [22, 65, 89] and genome annotation [90]. ProteomeLM demonstrates the power of integrating whole-proteome information across species. ProteomeLM reasons on proteomes at a coarse-grained level, representing each protein by a global representation from the protein language model ESM-C. These light representations allowed us to maintain reasonable context sizes and to work with full proteomes. In the future, the progress of long-context language models may enable ProteomeLM to directly operate at the amino-acid level across whole proteomes, paving the way to localized cross-protein interactions and more granular modeling of functional dependencies. One could even envision using full genomic nucleotide sequences, in the spirit of recent models that work on local genomic neighborhoods [17–19, 24–26]. While this may become possible, we expect models working on complete raw genomic sequences to be much heavier and to require much more training data, as they need e.g. to learn about coding regions and codons. Note that alternative architectures could help address longer sequences, as for protein language models [91, 92]. Models operating on raw genomic sequences can address regulatory functions or the role of non-coding DNA [17–19], which is beyond the scope of ProteomeLM. However, once they reach whole genomes, it remains to be seen whether they can efficiently and accurately predict proteome-level tasks such as PPI and gene essentiality, where ProteomeLM excels.

A major asset of ProteomeLM, allowed by our introduction of functional encodings instead of positional encodings, is that it can reason on proteomes across the tree of life. Nevertheless, in terms of absolute scores, ProteomeLM’s performance remains stronger on prokaryotes than on eukaryotes. This may be due to the relative scarcity of high-quality eukaryotic proteomes in the training dataset and to the higher complexity of eukaryotic genome organization and gene regulation. While fine-tuning ProteomeLM on eukaryotic-only data, or training it only on that data, brings minor improvements on eukaryotes, it comes at the cost of reduced performance on bacteria. As eukaryotic genomic databases continue to grow, larger training sets may allow us to overcome this limitation. Along these lines, expanding the training set to include more eukaryotic proteomes, including less-well annotated ones, and metagenomic proteomes, should further improve ProteomeLM’s performance across taxonomic groups.

Another interesting perspective for future work is to train ProteomeLM using embeddings from protein language models trained over additional modalities, beyond sequences, such as structure (SaProt [93], ProstT5 [94], ESM-3 [95], ProtTrek [96]). Exploiting the complementary information available in structure and sequences [97] may help ProteomeLM infer more complex functional relationships. This direction is particularly promising for PPI prediction, as shown by recent work [27, 98–102].

ProteomeLM opens the way to several applications. Given its computational efficiency at inference, it can be used to map functional networks and co-expression clusters, predict protein complex membership across species, and study the evolution of these systems. For the determination of direct physical interactions, it can serve as a fast and accurate screening step to prioritize candidate protein pairs for computationally intensive analyses, including structural prediction approaches such as Boltz [103, 104] and AlphaFold3 [105], or interface prediction frameworks such as PIONEER 2.0 [102]. Beyond interaction prediction, ProteomeLM enables high-throughput *in silico* screens of gene essentiality and comparisons across species, and could be extended to investigate joint essentiality [106–110] and environment-dependent essentiality [111]. ProteomeLM embeddings can further serve as a basis for other downstream tasks such as context-aware fitness prediction, opening the possibility of large-scale investigations of fitness landscapes.

## Methods

### Architecture and training of ProteomeLM

#### Dataset

We collected 31,947 proteomes from OrthoDB (version 12) [67]. Each of these proteomes is a list of protein sequences annotated by their respective orthologous groups. An orthologous group comprises descendants of a common ancestral gene, separated by speciation, and usually retaining the same function. It is linked to functional annotations from Gene Ontology (GO) [112, 113], describing protein localization and biological processes. However, here, we do not use these functional annotations. Our functional encoding (described below) only relies on orthologous groups. Note that the definition of an orthologous group operates at a specific level of orthology. Here, we use all these levels (see below).

#### Protein representations

We used the ESM-Cambrian (ESM-C) model with 600 million parameters [12] to represent each of the 162 million proteins in our dataset. ESM-C is a protein language model, trained on a vast corpus of sequences, which is a successor of ESM-2 [10]. Note that, contrary to ESM-3, which is multimodal [95], it is a sequence-only model. We selected ESM-C because of its strong performance in capturing the structural and functional properties of protein sequences. Each protein sequence was encoded by ESM-C, and the per-amino acid ESM-C embeddings were averaged to obtain a global embedding of the protein with a fixed dimension of 1152.

To optimize computational efficiency, protein sequences were batched based on their length, reducing the overhead associated with the model’s quadratic time complexity. The whole embedding computation process took 192 GPU-hours on one H100 GPU.

#### Functional encoding

In natural language processing [114], and also for protein sequences [9, 12, 115] and for genomic sequences [18, 22, 89], BERT models usually rely on positional encoding, which provides the model with information on the order of tokens (words in a sentence, amino acids in a protein chain, nucleotides along the genome). However, such a positional encoding is not appropriate for our purpose, given the lack of conservation of genomic order across diverse species, and the lack of correlation between proximity along the genome and functional relationship or interaction in eukaryotic genomes. Instead, we designed a *functional encoding* based on OrthoDB orthologous groups, which provides the model with information about the functional identity of each protein.

OrthoDB organizes orthologous groups in a tree that mirrors the taxonomy of life. At the finest level, a leaf orthologous group contains a set of orthologous proteins from closely related species (e.g., Hominidae). Each leaf group is nested within progressively broader groups defined at higher taxonomic levels (e.g., Mammalia, Vertebrata, Metazoa), up to a root encompassing all cellular organisms. Moving up this hierarchy, orthologous groups become more inclusive, reflecting more ancient and broadly conserved functions. We exploit this hierarchy to construct a multi-scale functional representation for each protein, as follows. First, for each leaf orthologous group, containing individual proteins but no subgroups, we average the ESM-C embeddings of all proteins in that group, yielding a vector characterizing that specific orthologous group. Second, we propagate these representations through the hierarchy. Going from leaves to root, we compute a vector for each parent group as the equal-weight average of its immediate children’s vectors. Having each child orthologous group contribute equally to the parent representation prevents large groups from dominating. Thus, every node in the OrthoDB hierarchy is assigned a vector. For each protein, we collect all vectors representing each group along the ancestral path from its leaf orthologous group to the root. This yields a set of vectors that describe the protein’s functional identity at each taxonomic level. Finally, for proteins which are not members of an OrthoDB group, we use their ESM-C embedding as their functional encoding.

During training, for each protein in a proteome, the functional encoding is obtained by randomly sampling (at each epoch) one vector from its ancestral path. This stochastic sampling exposes the model to functional descriptions at varying levels of specificity during training, encouraging it to learn relationships between proteins that hold at multiple evolutionary scales.

We find that these functional encodings outperform simpler ones, based on a discrete representation of orthologous group identities at the broadest taxonomic level (see Supplementary Section 6). Our use of hierarchically averaged ESM-C embeddings captures richer signal, especially for predicting physical PPI.

#### Architecture

We trained a transformer encoder from scratch to learn complex relationships between the ESM-C embeddings of different proteins in a proteome. The core of the model is the DistillBERT architecture, available in Hugging Face’s `transformers` library [116]. We used FlashAttention-2 [117] to accelerate training and inference.

Each proteome is represented as a list of protein embeddings with their associated functional encodings. For each protein, the ESM-C embedding and the functional encoding are given as inputs to ProteomeLM, through two separate linear projection layers.

#### Training objective and loss design

ProteomeLM is trained using a masked language modeling (MLM) objective adapted to proteome-level inputs. During the training of ProteomeLM, we limit the size of the proteomes to 4096 by randomly subsampling proteins when proteomes are longer. While longer inputs could in principle be employed, since we use FlashAttention, which has linear memory complexity, we chose to limit the length of the input to 4096 to reduce computational time, which still has quadratic complexity. We randomly mask 50% of the protein representations within a proteome, while their functional encodings are kept unmasked. Masked proteins are replaced by their functional encoding, and the model is trained to reconstruct the original protein embeddings based on contextual signals from the rest of the proteome.

The standard masked language modeling loss cannot be applied here, because we work directly with continuous embeddings and not with discrete tokens. A straightforward alternative would be the mean squared error (MSE) between the actual embedding *x* and the predicted embedding *x̂*. However, this approach results in a degenerate solution where the model simply reproduces the functional encoding *x̅*. Indeed, when expressed in terms of the residuals *r* = *x* − *x̅* and *r̂* = *x̂* − *x̅*, the MSE becomes insensitive to the angle *θ* between *r* and *r̂* when *x* approaches *x̅*, and thus fails to penalize incorrect residual directions (see Supplementary Section 7). To address this challenge, we decouple the residual magnitude and direction. We formulate the residual prediction problem probabilistically under this decoupling, using tractable assumptions on the distributions of residual magnitudes and angles. A maximum-likelihood derivation under this model leads to the following custom polar loss (see Supplementary Section 7):

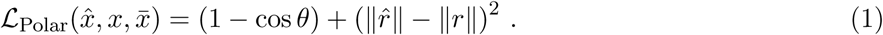

The first term corresponds to the cosine embedding loss applied to the residuals, and encourages alignment between *r* and *r̂*. The second term penalizes discrepancies in residual magnitude through the squared difference between the Euclidean norms of *r* and *r̂*. The polar loss is minimized if and only if *x̂* = *x*, thereby avoiding the failure mode of the MSE loss. Empirical comparisons are given below, in the paragraph titled “Comparison of losses”.

#### Training dynamics and scaling behavior

We trained four variants of ProteomeLM, differing by model sizes: XS (5.6M parameters), S (36M), M (112M), and L (328M). All models were trained for 210 epochs on a dataset comprising 31,000 proteomes with a total of 160 million proteins. Validation loss was measured on a 2% held-out set of proteomes randomly sampled from the training set.

Training remained stable across all model sizes, showing smooth convergence, see Figure 6A. Figure 6A and B show that performance, assessed by loss value, improved steadily from XS to M, suggesting that the model benefits from increased capacity. However, the L model failed to outperform M, and in some cases showed degraded performance. In particular, in Figure 6B, the trend follows a scaling law from XS to M, before performance degrades for the L model. We attribute this to overfitting, given that the number of trainable parameters exceeds the number of unique training proteins in the training set for ProteomeLM-L. To rule out architecture-specific factors, we tested variants of ProteomeLM-L with different numbers of layers, heads, and embedding dimensions. These variants exhibited similar behavior in early training, reinforcing the interpretation that training data volume is the limiting factor.

**Figure 6:**
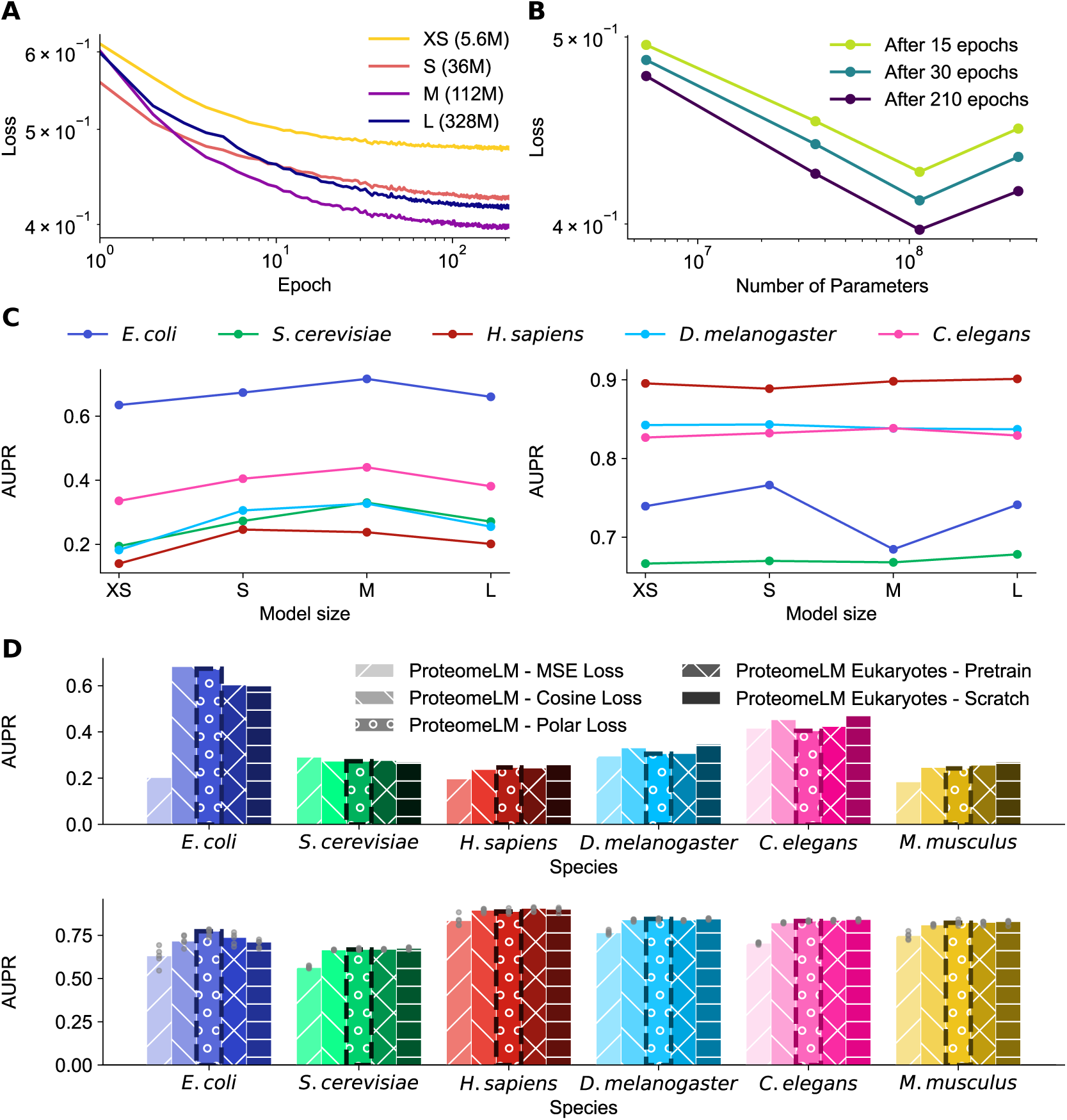
Training dynamics, scaling behavior, and training strategies for ProteomeLM. **(A)** Evaluation loss during training for four ProteomeLM model sizes: XS (5.6M parameters), S (36M), M (112M), and L (328M). **(B)** The final evaluation loss is shown for three different epochs. **(C)** The AUPR for unsupervised PPI prediction using summed attention coefficients over all heads and layers (left) and supervised PPI prediction through the ProteomeLM-PPI architecture (right) is shown across five species on the D-SCRIPT dataset. **(D)** Comparison of three different loss functions (MSE, cosine and polar, where polar is the one retained throughout), and evaluation of two training strategies focused on eukaryotic data (fine-tuning and training from scratch). AUPR are shown for both unsupervised (top) and supervised (bottom) PPI prediction tasks. As in C (left panel), summed attention coefficients over all heads and layers are used for unsupervised prediction. C-D: All models used in these comparisons were trained for 210 epochs under identical hardware and random seed conditions to ensure fair evaluation.

In Figure S11, we show the training dynamics of each attention head in ProteomeLM-S by tracking their AUC for unsupervised PPI recovery during training. Certain heads become increasingly predictive of PPI as training progresses. Some of them display taxon-specific specialization, while others exhibit consistent predictive power across species. These results highlight that ProteomeLM’s attention heads learn distinct, biologically meaningful signals during training.

We also used the D-SCRIPT dataset [27] to assess the performance of the models on PPI recovery (training and validation on disjoint sets of *H. sapiens* PPI, test on other species). The left panel of Figure 6C shows that, for unsupervised PPI recovery, AUPR increases with model size up to M, and decreases for size L, thus confirming the trend observed for the loss. Note also that performance on human data slightly decreased from S to M, indicating that larger models do not always generalize better across all species. The right panel of Figure 6C shows that AUPR increases over model size on eukaryotes, while it degrades after model S on *E. coli*, again showing the better generalization capabilities of the smaller models.

#### Comparison of losses

We evaluated our polar loss function against two natural alternatives, namely the mean squared error (MSE) loss between *x̂* and *x* and the cosine embedding loss between *x̂* − *x̅* and *x* − *x̅*. (Note that we do not consider the cosine embedding loss between *x̂* and *x* because these vectors tend to have a similar orientation anyway within each protein family.) For our comparison of loss functions, we trained a version of ProteomeLM for 72h with each of these two alternative losses. Performance comparisons were conducted in both unsupervised and supervised protein-protein interaction (PPI) prediction tasks. In the unsupervised setting, we used the mean of attention weights over all heads and layers as a simple estimate of interaction probability for each protein pair. In the supervised setting, we trained a downstream classifier (see Figure 4A) using frozen representations obtained from models trained with each loss function.

As shown in Figure 6D, our polar loss consistently outperformed both the MSE loss and the cosine loss across multiple evaluation metrics. In particular, the unsupervised AUC scores were generally performing higher when the model was trained with the polar loss or the cosine embedding loss than with the MSE loss. Likewise, in the supervised PPI prediction task, the models trained with the polar loss yielded higher precision-recall performance (AUPR). The MSE loss showed the weakest performance in both settings. As mentioned above, it is because the model then converges towards a degenerate solution, where the model simply reproduced the functional encoding (*x̂* = *x̅*).

Note that, in addition to predictive accuracy, we observed that the polar loss produced embeddings with properties more closely aligned to those of ESM-C, thus facilitating interoperability between models, e.g. improving compatibility in transfer learning applications. These results validate the polar loss as an appropriate objective for reconstructing protein embeddings within a proteome context.

#### Improving performance on eukaryotic data

We observe that ProteomeLM’s accuracy is comparatively lower on eukaryotic datasets than on prokaryotic ones (see Figures 2 and 4). We explored two approaches to improve performance on eukaryotes: fine-tuning the pretrained ProteomeLM on eukaryotic data, and training a new model from scratch on eukaryotic data only.

Figure 6D shows that both approaches provide moderate improvements on eukaryotic benchmarks, both for unsupervised and supervised PPI prediction. However, these improvements come with a decrease in performance on prokaryotes such as *E. coli*, indicating a trade-off between specialization and generalization.

Given our goal to design a model that works across diverse organisms, we retained the baseline ProteomeLM models for our main analyses. However, the specialized eukaryotic alternatives can be valuable for specific applications.

### Supervised protein-protein interaction prediction: ProteomeLM-PPI

#### Architecture and input

The supervised ProteomeLM-PPI model relies on a modular neural network that processes both individual protein embeddings (node features) and attention coefficients (edge features) through distinct but integrated modules, see Figure 4A.

Specifically, the node feature module reduces each protein embedding from 640 to 256 dimensions through two layers (640 → 512 → 256), with layer normalization and dropout to enhance stability and weight regularization. To model the interaction between two proteins, the network combines their transformed representations by concatenating each of the two representations, their element-wise multiplication, and their absolute difference, thus resulting in a 1024-dimensional vector (4 × 256), which is then compressed to 64 dimensions through the interaction processor (1024 → 128 → 64). In parallel, the edge feature module reduces the 48-dimensional input (from the 48 attention heads of ProteomeLM-S) to 32 dimensions (48 → 64 → 32). The 64-dimensional processed interaction features from the interaction processor are then concatenated with the 32-dimensional pairwise feature vector to form a 96-dimensional input to the final PPI prediction classifier module. This input then passes through two layers (dimensions: 96 → 128 → 64), before the PPI predictor outputs a final interaction score. This modular design enables the model to flexibly integrate different sets of learned features while maintaining strong inductive biases for capturing protein-protein relationships.

#### Training

To train the model, we used a train-validation-test split. During training, the model is optimized on the training set, while its performance is monitored on the validation set to apply early stopping, preventing overfitting. During training, the proteins pairs of the training set are fed into the model in mini-batches as triplets comprising ProteomeLM embeddings of both proteins involved, and ProteomeLM attentions weights between the two of them. The training minimizes binary cross-entropy with logits using the Adam optimizer [118]. At each epoch, predictions on the validation set are evaluated using the area under the precision-recall curve (AUPR), and the best model state is saved based on this metric. After training, the best model is evaluated on the held-out test set, and we report AUC and AUPR.

Our training approach ensures a fair assessment of the model’s generalization ability by performing model selection on a dedicated validation set, rather than the test set, thereby avoiding overfitting to the test data. For both the dataset from [76] and the D-SCRIPT dataset [27], clean and non-overlapping splits into training, validation, and test sets were already available, and we employed them.

### Supervised gene essentiality prediction: ProteomeLM-Ess

#### Architecture and input

ProteomeLM-Ess is a two-layer fully connected classifier that takes as input embeddings from any ProteomeLM model, has a hidden layer of size 2048, and outputs two logits, which are normalized with a softmax function to obtain an essentiality score. In the hidden layer, ProteomeLM-Ess has a ReLU activation and dropout with probability 0.5. Protein embeddings are normalized using the genome-wide mean and standard deviation before being given as input to ProteomeLM-Ess.

#### Data and training

ProteomeLM-Ess is trained in a supervised way, using ProteomeLM embeddings together with essentiality labels from the OGEE database [77]. We recovered the protein sequences associated to the essentiality labels from other databases, by matching the gene names (gene IDs) provided by OGEE. Specifically, we collected protein sequences from UniProt [119], NCBI [120], *Saccharomyces* Genome Database (SGD) [121] and Fitness Browser [122], and obtained data for 87 taxonomic IDs. Relying on curated complete proteomes whenever possible allowed to minimize ambiguities coming from the presence of isoforms or duplicate protein sequences (i.e. identical sequences with different protein IDs). The remaining duplicate entries were merged, while keeping both IDs.

Out of the 87 total genomes, we used 83 to train ProteomeLM-Ess, holding out the genomes of *S. cerevisiae* and of 4 strains of *E. coli*. We also collected essentiality data for the synthetic cells JCVI-Syn1.0 [78] and JCVI-Syn3A [79, 80], to evaluate the model after training. For the 83 genomes used for training, we split the proteins into training, validation and test sets by clustering proteins across all genomes according to sequence similarity. Specifically, we clustered sequences using MMSeqs2 [123] with a 40% similarity threshold. We designed our split so that if two labeled proteins belong to the same cluster, then they are either both in the training set, in the validation set, or in the test set. The data split is performed at the protein level and not at the genome level, to avoid the model relying on sequence similarity between, say, two orthologs in similar genomes, as a shortcut to predict essentiality. All protein sequences are given as input to ProteomeLM to build contextualized embeddings. The training procedure and objective used for ProteomeLM-Ess are the same as the ones used for ProteomeLM-PPI (see above).

## Data and code availability

We provide code for training and using ProteomeLM at https://github.com/Bitbol-Lab/ProteomeLM. Code weights, and all the material needed to reproduce our results, are available there.

## Acknowledgments

This research was funded by the European Research Council (ERC) under the European Union’s Horizon 2020 research and innovation programme (grant agreement No. 851173, to A.-F. B.). We thank Jing Zhang and Ian R. Humphreys for useful discussions and additional material for PPI screening, Liedewij Laan for useful discussions about gene essentiality and James Sáenz for useful discussions about minimal cells.

## Supplementary Information

### 1 Distinguishing physical interactions from functional associations

ProteomeLM learns statistical dependencies between proteins based on their context across thousands of genomes, and shares similarities with phylogenetic profiling [49]. Do ProteomeLM’s attention coefficients capture direct physical binding, broader functional associations, or both? To address this, we compare ProteomeLM attention coefficients across different types of protein-protein associations.

#### Benchmark construction

For *E. coli*, *S. cerevisiae*, and *H. sapiens*, we constructed a benchmark comprising the following four categories of protein pairs:

- **Direct interactions (PDB):** Protein pairs involved in the same PDB complex, with buried surface area above 500 Å^2^ and at least 10 pairs of residues with distance between closest atoms below 8 Å.
- **Same complex interactions (PDB):** Protein pairs involved in the same PDB complex but that do not satisfy the criteria stated above.
- **Coexpression (STRING):** Protein pairs with high coexpression scores (≥ 0.7) in the STRING database [69], representing proteins with correlated expression patterns across experimental conditions.
- **Random pairs:** Randomly sampled protein pairs from the same proteome, serving as negative controls. We ensured that every protein in these pairs is involved in known interactions (from the three previous sets).

Pairs were filtered to ensure no overlap between categories, and random pairs were sampled to exclude any pair present in STRING with non-zero scores.

#### Results

We first evaluate the strength of the signal in ProteomeLM attention coefficients for each interaction type. Table S1 shows the Area Under the Receiver Operating Characteristic curve (AUROC) for discriminating each interaction type from random pairs, using the mean of ProteomeLM attention coefficients from all heads and layers. Across all three species, coexpression pairs yield the strongest attention signals, achieving AUROC values of 0.92-0.95. Direct and physical interactions are also well detected (0.75-0.92), but with a lower strength than coexpression. Hence, ProteomeLM is a powerful predictor of functional associations and gene co-regulation.

**Table S1:** Discriminating three interaction types from random pairs. All results are given in terms of AUROC. Higher values indicate a stronger attention signal relative to a random background.

| Species | Direct | Complex | Coexpression |
| --- | --- | --- | --- |
| *E. coli* | 0.873 | 0.916 | 0.924 |
| *S. cerevisiae* | 0.810 | 0.828 | 0.946 |
| *H. sapiens* | 0.745 | 0.787 | 0.946 |

Next, to assess whether ProteomeLM can distinguish physical binding from broader functional associations, we train logistic regression classifiers to discriminate between interaction types based on attention coefficients. Table S2 demonstrates that coexpression can be well distinguished from direct or same-complex interactions using ProteomeLM. For instance, the classification accuracy when distinguishing direct interactions from coexpression has a 0.89 AUROC in *S. cerevisiae*. However, the more subtle distinction between direct and same-complex interactions is less well captured by ProteomeLM.

**Table S2:** Pairwise binary classification between interaction types. Ability of logistic regression classifiers trained on ProteomeLM attention heads to distinguish between specific interaction types. All results are given in terms of AUROC.

| Species | Direct vs. complex | Direct vs. coexpression | Complex vs. coexpression |
| --- | --- | --- | --- |
| *E. coli* | 0.504 | 0.728 | 0.626 |
| *S. cerevisiae* | 0.552 | 0.885 | 0.851 |
| *H. sapiens* | 0.601 | 0.904 | 0.882 |

#### Interpretation

These results demonstrate that ProteomeLM is an excellent predictor of broad functional relationships, and also captures specific information regarding physical binding. Furthermore, the attention signatures for physical and genetic interactions are distinct. This suggests a dual utility: ProteomeLM can be used to map functional networks and co-expression clusters, while simultaneously serving as a high-recall filter for structural PPI prediction pipelines by identifying pairs that are physically interacting.

### 2 Application of ProteomeLM to structurally resolved protein complexes

#### 2.1 *E. coli* ribosome

To assess ProteomeLM’s ability to recover physical protein interactions within a multi-protein complex, we consider the *E. coli* 70S ribosome, focusing on ground-truth inter-chain contacts extracted from a high-resolution cryo-EM structure (PDB: 7K00, 2.0 Å resolution) [124]. This complex comprises 46 proteins, and provides a benchmark involving a dense network of both direct and indirect PPI within a complex.

##### Ground truth extraction

We parsed the atomic coordinates of the structure and mapped each protein chain to its UniProt identifier in the *E. coli* K-12 proteome. For each pair of ribosomal proteins, we computed the minimum C*α*-C*α* distance. We define direct interactions as protein pairs where this distance is below 8 Å, yielding a sparse ground-truth direct interaction matrix among the 46 ribosomal proteins.

##### Complex membership detection

We first ask whether ProteomeLM attention scores distinguish ribosomal proteins from the rest of the proteome. For each pair of ribosomal proteins (intra-complex) and for pairs consisting of one ribosomal and one non-ribosomal protein (inter-complex), we compute the mean attention score across all heads and layers. As shown in Figure S1A, intra-complex pairs receive significantly higher attention scores than inter-complex pairs. Quantitatively, ProteomeLM achieves an AUROC of 0.993 for distinguishing intra- from inter-complex pairs, confirming that the model reliably identifies co-complex membership at the proteome scale.

**Figure S1:**
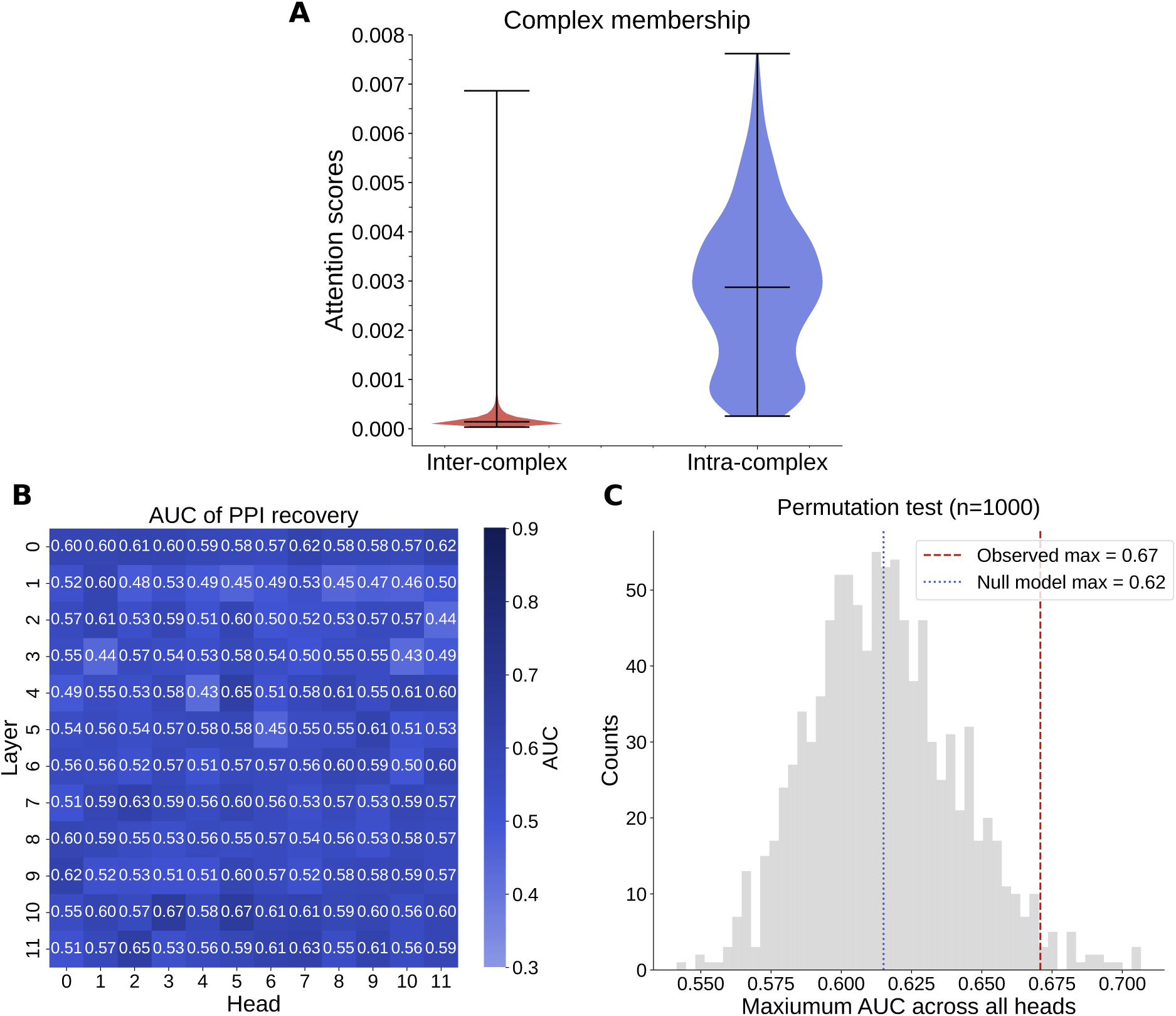
Validation of ProteomeLM on the *E. coli* 70S ribosome (PDB: 7K00). **(A)** Distribution (with extremes and median) of ProteomeLM attention scores for intra-complex (ribosomal–ribosomal) versus inter-complex (ribosomal–non-ribosomal) protein pairs. Intra-complex pairs receive substantially higher attention (membership AUROC: 0.993). **(B)** Per-head AUROC for predicting ground-truth direct interactions (minimum C*α*-C*α* distance *<* 8 Å) among the 46 mapped ribosomal proteins, for each attention head of ProteomeLM-M (12 layers, 12 heads). The best head achieves an AUROC of 0.671. **(C)** Permutation test (*n* = 1,000) for the maximum AUROC across all heads. The observed maximum (red dashed line, 0.671) significantly exceeds the expected null maximum (blue dotted line, 0.615), with a p-value of 0.024.

##### Direct interaction prediction

We next ask whether ProteomeLM attention coefficients can recover protein pairs in direct interaction *within* the ribosome. We evaluate the AUROC of each individual attention head for predicting the ground-truth direct interaction matrix among the 46 ribosomal proteins (Figure S1B). The best-performing head yields an AUROC of 0.671. To assess statistical significance, we perform a permutation test. Specifically, we use 1,000 permutations of the direct interaction labels, and consider the distribution of maximum AUROC across all heads in this case as the null distribution ( Figure S1C). The observed maximum significantly exceeds the expected null maximum (0.615), with a p-value of 0.024, indicating that ProteomeLM encodes a statistically significant structural signal within this complex. However, signal is less strong than for complex membership detection (in line with Section 1). We further observe a weak but significant negative correlation between attention scores and physical distance among ribosomal protein pairs (Pearson correlation: −0.22), indicating that spatially close proteins tend to feature higher attention.

##### Interpretation

These results are consistent with the design and training objective of ProteomeLM. As a proteome-scale model trained to predict masked protein embeddings from genomic context, ProteomeLM learns functional co-dependencies between proteins, rather than direct interactions. The near-perfect complex membership detection (AUROC of 0.993) demonstrates that ProteomeLM robustly captures which proteins belong to the same complex, confirming the results of Section 1 on a specific example. The weaker but significant intra-complex contact signal suggests that ProteomeLM encodes some information about the internal organization of complexes, and that there is room for improvement. Recall that ProteomeLM operates on whole-protein embeddings and does not have access to residue-level information. ProteomeLM can be employed as a fast and precise filter to identify candidate PPI pairs to be studied by more detailed but computationally much heavier models, such as Boltz [103, 104], or AlphaFold3 [105].

#### 2.2 *S. cerevisiae* TRiC/CCT chaperonin

As another test case, we consider the eukaryotic TRiC/CCT chaperonin, which is a complex of 8 paralogous subunits (CCT1-8) arranged in a specific circular order [125]:

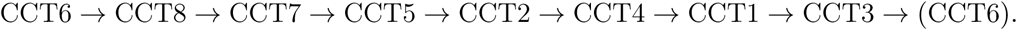

This provides an interesting and challenging benchmark, because all 8 subunits are paralogs with high sequence similarity, all belong to the same complex, but only 8 out of 28 possible subunit pairs are in direct interaction within the ring. We processed the *S. cerevisiae* proteome through ProteomeLM and extracted attention coefficients for all possible pairs of CCT subunits.

##### Complex membership detection

We first ask whether ProteomeLM attention scores distinguish CCT complex members from non-members, as for the ribosome (Section 2.1). For each intra-complex pair (CCT-CCT, 28 pairs) and inter-complex pair (CCT-non-CCT, 8,000 pairs using 1,000 non-CCT proteins sampled uniformly at random), we compute the mean attention score across all heads and layers. As shown in Figure S2A, intra-complex pairs receive higher attention than inter-complex pairs, achieving a perfect membership classification. This confirms that ProteomeLM reliably identifies co-complex membership for this complex of paralogs, consistent with the results obtained in Sections 1 and 2.1.

##### Ring arrangement recovery

We next ask whether ProteomeLM can resolve the specific circular arrangement of subunits within the ring. Specifically, we evaluate whether individual attention heads discriminate adjacent from non-adjacent subunit pairs. The best-performing head achieves an AUROC of 0.731 ( Figure S2B). However, a permutation test shows that this is not significant. We further enumerate all 5,040 circular permutations of the 8 subunits and score each by the total attention between consecutive pairs. The experimentally determined ring order [125] ranks 305th out of 5,040 (top 6.1%, *z* = 1.58), which does not reach statistical significance at conventional thresholds.

**Figure S2:**
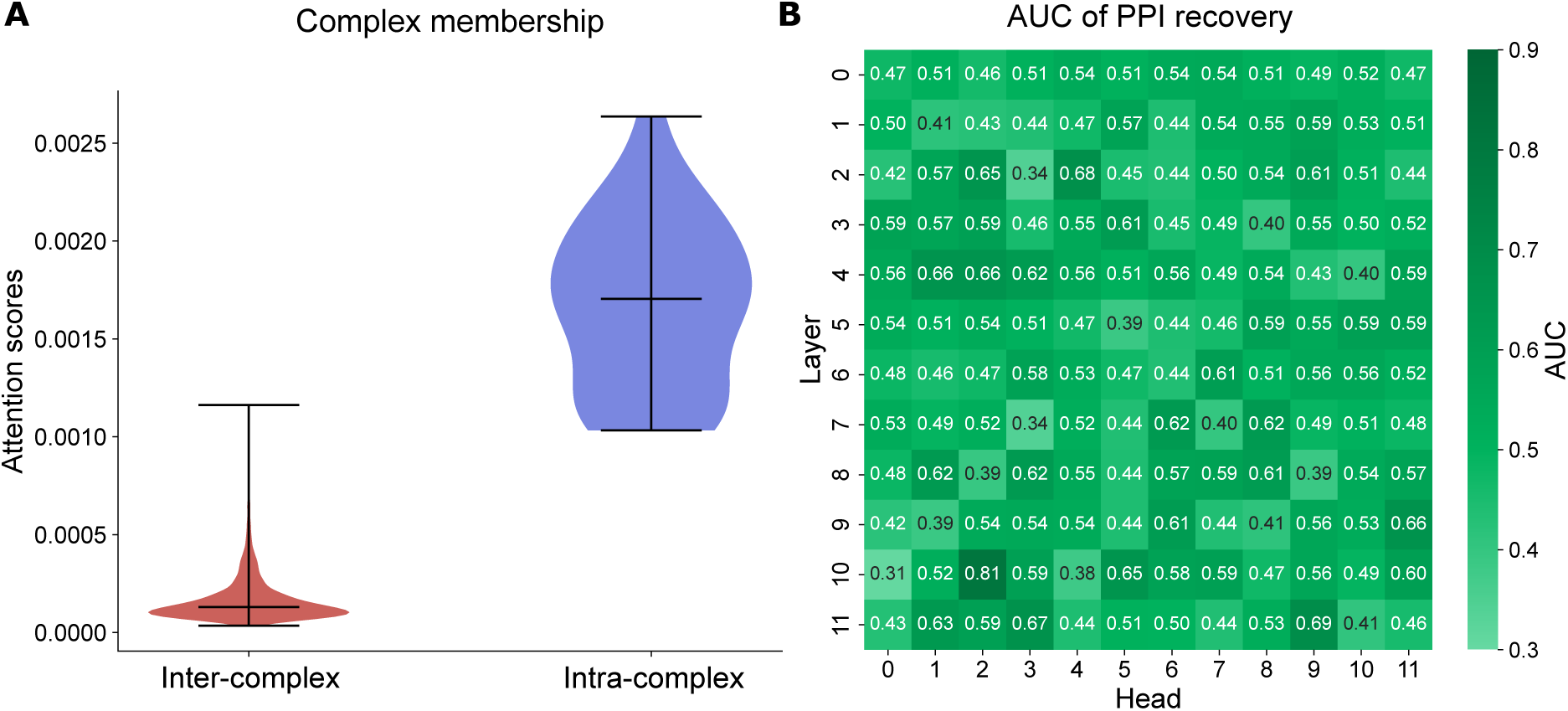
ProteomeLM identifies TRiC/CCT complex membership but does not resolve the ring arrangement. **(A)** Distribution of ProteomeLM attention scores for intra-complex (CCT–CCT) and inter-complex (CCT–non-CCT) protein pairs. Intra-complex pairs receive on average 8.4 times higher attention (Mann-Whitney U test p-value: 2.93 × 10*^−^*^20^; membership AUROC = 1.00). **(B)** Per-head AUROC for discriminating adjacent from non-adjacent subunit pairs (8 adjacent out of 28 total), for each attention head of ProteomeLM-M (12 layers, 12 heads). The best head achieves an AUROC of 0.731, but a permutation test (*n* = 1,000) shows that this falls below the expected null maximum (0.793; p-value: 0.911), indicating no significant adjacent-pair signal.

##### Interpretation

These results corroborate those obtained in Section 1: ProteomeLM reliably identifies which proteins belong to the same functional complex, but does not resolve fine-grained subunit arrangement within a complex as well. For TRiC/CCT, the 8 subunits descend from a common ancestor and occupy structurally equivalent positions in the ring. Hence, from the perspective of evolutionary co-occurrence patterns, which ProteomeLM captures through proteome-level context, they are nearly interchangeable. The identification of direct PPI interactions is thus expected to be more challenging here than in the *E. coli* ribosome studied in Section 2.1, in line with our results.

### 3 Improvement of learned representations over input representations

ProteomeLM takes as input per-protein embeddings from ESM-C, which already encode evolutionary and functional information. An important question is whether ProteomeLM’s attention coefficients simply encode similarities already present in the input embeddings, or whether they extract a more subtle signal. Furthermore, the functional encodings employed in ProteomeLM may encode further evolutionary information, since their construction involves a hierarchical averaging within orthology groups. This could potentially be relevant for the prediction of PPI, e.g. because interacting proteins have similar phylogenies [126, 127]. Here, we address these two points through comparative analyses.

#### 3.1 Cosine similarity of input embeddings and functional encodings

For each species (*E. coli*, *S. cerevisiae*, *H. sapiens*), we compute the all-versus-all cosine similarity between ESM-C input embeddings and evaluate its ability to discriminate interacting from non-interacting pairs across three interaction types (direct, same complex, coexpression), using the benchmark described in Section 1. As shown in Table S3, ESM-C embeddings already capture information about PPI, achieving AUROC values of 0.61-0.75 for direct interactions and 0.74-0.83 for coexpression. In addition, we compute the all-versus-all cosine similarity between functional encodings. We obtain a weaker signal, with AUROC 0.58-0.70 for direct interaction and 0.46-0.68 for coexpression.

ProteomeLM’s attention coefficients consistently and substantially outperform the baseline provided by cosine similarity between ESM-C embeddings. Specifically, the best individual ProteomeLM attention head achieves AUROC improvements of 0.09-0.16 over cosine similarity for direct interactions and 0.09-0.18 for coexpression (Table S3). Many attention heads outperform the ESM-C cosine similarity baseline (see Figure S3A), indicating that ProteomeLM learns nontrivial PPI-relevant information in its attention heads.

**Table S3:** PPI prediction using cosine similarity of input embeddings, of functional encodings, and best ProteomeLM attention head. Results are reported across three interaction types (see Section 1) in three species. All results are given in terms of AUROC.

| Species | Method | Direct | Complex | Coexpression |
| --- | --- | --- | --- | --- |
| *E. coli* | Input embedding | 0.753 | 0.745 | 0.741 |
|  | Functional encoding | 0.702 | 0.660 | 0.655 |
|  | Best ProteomeLM head | 0.866 | 0.887 | 0.918 |
| *S. cerevisiae* | Input embedding | 0.688 | 0.671 | 0.782 |
|  | Functional encoding | 0.642 | 0.612 | 0.680 |
|  | Best ProteomeLM head | 0.782 | 0.826 | 0.942 |
| *H. sapiens* | Input embedding | 0.606 | 0.619 | 0.830 |
|  | Functional encoding | 0.584 | 0.556 | 0.461 |
|  | Best ProteomeLM head | 0.721 | 0.745 | 0.946 |

#### 3.2 PCA removal from the input similarity matrix

We perform a PCA decomposition of the all-versus-all cosine similarity matrix between ESM-C input embeddings, as well as of the one between functional encodings. In each of these two cases, we progressively project out the top *k* principal components (for *k* = 0, 1, 5, 10, 20, …, 148), and we use the residual similarity scores to predict PPIs. As shown in Figure S3A, PPI prediction performance decreases rapidly with PC removal, falling below the original cosine baselines. This indicates that the dominant modes of pairwise similarity in the input space carry PPI-relevant signal. However, that signal is weaker than the one encoded in ProteomeLM attention heads. Thus, ProteomeLM’s attention does not operate by simply filtering out top components of the input similarity matrix. We also note that the signal degrades more slowly in the functional encoding decomposition than in the input embedding decomposition.

#### 3.3 Supervised evaluation with sequence-similarity-controlled splits

To further validate that ProteomeLM learns interaction-specific representations beyond sequence similarity, we constructed rigorous supervised benchmarks for three species using high-confidence interactions from PDB [70]. We applied sequence-similarity-aware train/validation/test splits: proteins were clustered at 40% sequence identity using MMseqs2 [123], and entire clusters were assigned to train/validation/test (50/20/30) via min-cut optimization [128] that minimizes lost cross-split interactions. This ensures that no protein in the test set shares more than 40% sequence identity with any protein seen during training, thereby preventing homology-based information leakage.

We train simple multilayer perceptron classifiers (comprising one hidden layer with 64 units, and using ReLU activation functions) on this data, using different feature combinations:

- ESM-C embeddings alone (concatenated per-protein representations);
- ProteomeLM attention coefficients alone;
- ProteomeLM embeddings alone;
- ProteomeLM embeddings and attention coefficients.

**Figure S3:**
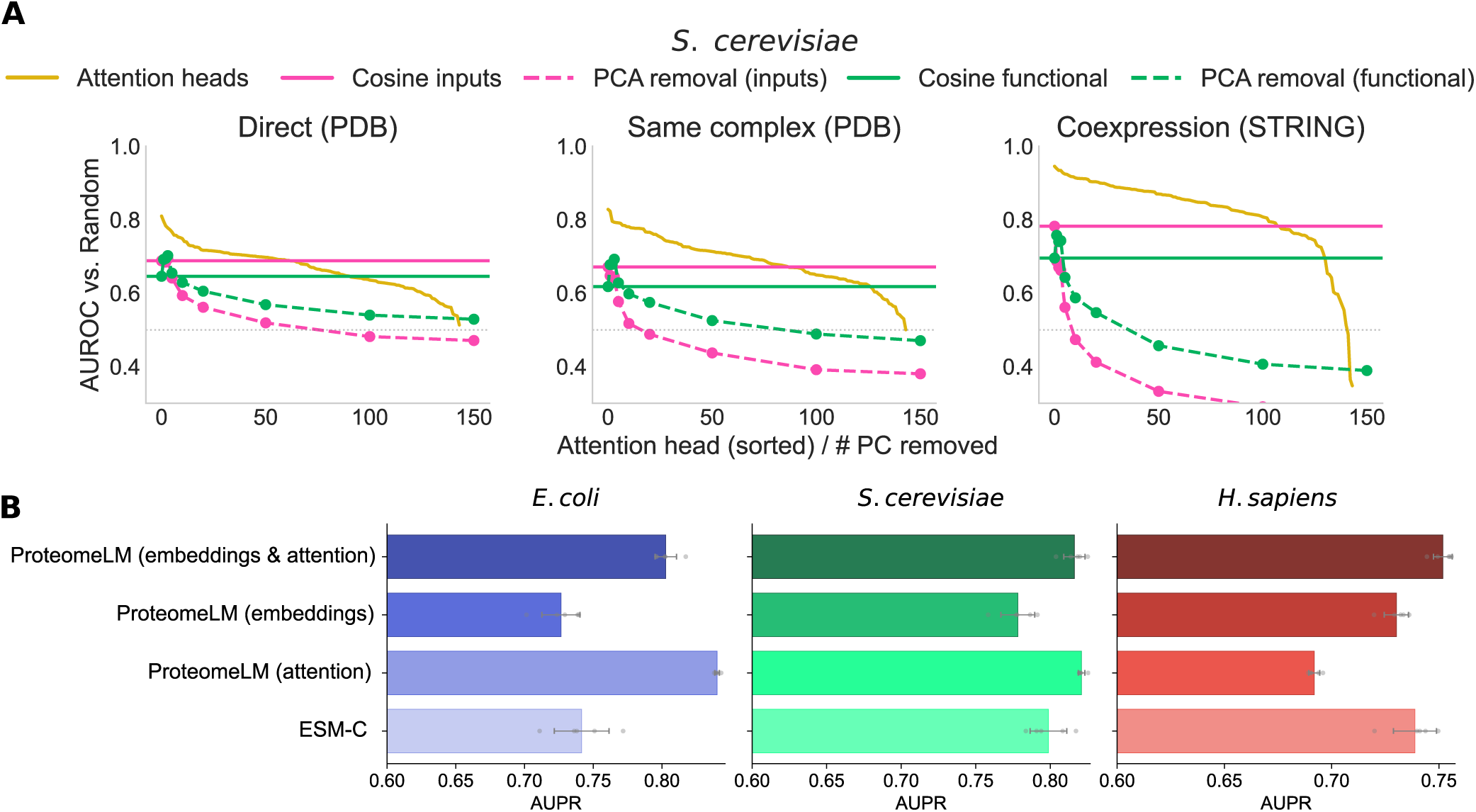
ProteomeLM representations improve over input embeddings for PPI prediction. **(A)** For each interaction type (direct, same complex, coexpression), we compare three unsupervised approaches: individual ProteomeLM-M attention heads sorted by AUROC, cosine similarity of ESM-C input embeddings, and cosine similarity of ProteomeLM functional encodings. For the latter two, we also perform progressive removal of the top *k* principal components. **(B)** AUPR of supervised multilayer perceptron classifiers trained on four feature sets, using ProteomeLM-M under sequence-similarity-controlled splits (40% identity): ESM-C embeddings alone, ProteomeLM attention alone, ProteomeLM embeddings alone, and the combination of ProteomeLM embeddings and attention. Error bars: standard deviation over 5 replicates (each shown with a gray marker).

As shown in Figure S3B, ProteomeLM attention coefficients alone outperform ESM-C embeddings in *E. coli* (AUPR of 0.80 vs. 0.74), and perform comparably in *S. cerevisiae*. Furthermore, the combination of ProteomeLM embeddings and attention coefficients consistently yields better performance than ESM-C embeddings across all three species. This result demonstrates the benefit of ProteomeLM features over the ESM-C baseline. It also confirms that ProteomeLM embeddings and attention coefficients carry complementary information. As elsewhere, we note that ProteomeLM performs particularly well on *E. coli*.

### 4 Comparison of compute requirements for human interactome scanning

Here, we focus on the specific problem of scanning the whole human interactome. A recent large-scale study applied DCA systematically to over 190 million *Homo sapiens* protein pairs [72]. We estimate the floating-point operations (FLOPs) required for this, as well as for each stage of our own screening with ProteomeLM. For this, we employ the peak FP32 throughput of relevant GPUs and the reported wall-clock runtimes. All values represent theoretical upper bounds. The results of the comparison below are summarized in Figure 3A.

#### DCA [72]

According to the authors of Ref. [72] (Jing Zhang, private communication), the Direct Coupling Analysis (DCA) step was performed over 1–2 months using between 50 and 100 GPUs, including NVIDIA RTX 6000, RTX 8000, A100, and A40. For our estimate, we assume:

- 75 concurrent GPUs over 45 days (i.e., 81,000 GPU-hours);
- Even distribution across the four GPU models;
- Peak FP32 throughput: 16.3 TFLOPS (RTX 6000 and RTX 8000), 19.5 TFLOPS (A100), and 37.4 TFLOPS (A40).

The resulting compute requirement is:

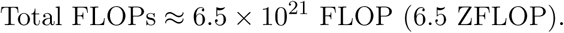

#### ProteomeLM training

Our model was trained on a single NVIDIA H100 SXM5 GPU during 72 hours. With a peak FP32 throughput of 67 TFLOPS, the total estimated compute is:

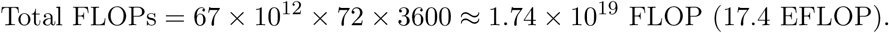

This value assumes uninterrupted training with full GPU utilization.

#### ProteomeLM inference

A 10-minute inference run was performed on an NVIDIA RTX A6000 GPU. With a peak FP32 throughput of 38.7 TFLOPS, the total compute estimated is:

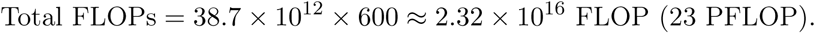

### 5 Comparison of ProteomeLM-Ess with other gene essentiality prediction methods

In Figure S4, we provide a comparison of ProteomeLM-Ess with different state-of-the-art gene essentiality prediction methods. During the training of ProteomeLM-Ess, *E. coli* and *S. cerevisiae* were held out, and hence, none of the gene essentiality labels of these species were seen. Among the alternative methods considered, some of them used part of the labeled essentiality data from the two species of interest for training (see the list below for details). We observe in Figure S4 that ProteomeLM-Ess outperforms all other methods considered on *E. coli*. For *S. cerevisiae*, ProteomeLM-Ess outperforms all three methods that did not include labeled data from that species in training, by a substantial margin. However, the methods that included such data in training perform as well as or better than ProteomeLM-Ess. This suggests that their training set gives them an intrinsic advantage with respect to ProteomeLM-Ess. Overall, ProteomeLM-Ess is state of the art for *E. coli*, and also for *S. cerevisiae* among methods that do not include labeled data from that species in the training set. Note also that, among all methods considered here, Evo [24] and Evo 2 [25] are the only fully unsupervised methods, which explains their lower performance.

Figure S4 compares ProteomeLM-Ess to the following methods on *E. coli* :

- **ZUPLS**: Song et al. [81] use a partial least squares algorithm trained on sequence-derived features. The AU-ROC value shown in Figure S4 is the one they report when predicting *E. coli* essential genes by training the model on *B. subtilis* essential genes.
- **Azhagesan et al.**: Azhagesan et al. [82] train a random forest classifier on data from 27 prokaryotic organisms, using a combination of features derived from the sequence and from the protein-protein interaction network. The value of the AU-ROC shown in Figure S4 is the one they report for the leave-one-species-out validation for *E. coli*, i.e. when training the model on data from all organisms except *E. coli*.

**Figure S4:**
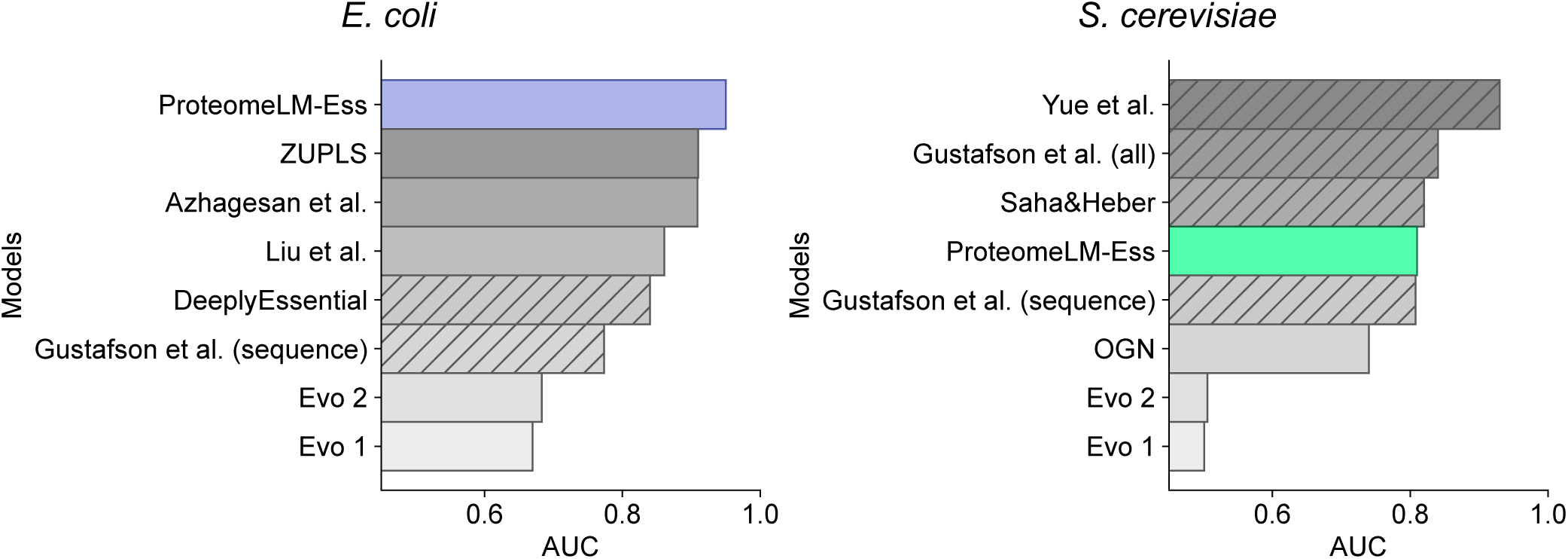
Performance of ProteomeLM-Ess and of other methods for gene essentiality prediction. The performance of ProteomeLM-Ess at predicting essential genes is compared to state-of-the-art methods for gene essentiality prediction [24, 25, 81–88], for *E. coli* (left) and *S. cerevisiae* (right). These other methods are briefly described in Section 5. Hatches indicate methods that use part of the organism’s genes as labeled training data, contrary to ProteomeLM-Ess, where *E. coli* and *S. cerevisiae* were held out from the training set.

- **Liu et al.**: Liu et al. [83] identify 40 sequence-derived features, and train a support vector machine classifier using data from 31 bacterial species. The AU-ROC value shown in Figure S4 is the one they report for the leave-one-species-out validation for *E. coli*.
- **DeeplyEssential**: Hasan and Lonardi [84] use data from 30 bacterial species and train a multilayer perceptron with 6 hidden layers using only sequence-derived features. The AU-ROC value shown in Figure S4 is an upper bound to the AU-ROC values they obtain for all Gram-negative bacteria. They aggregate data from all species and use 80% of genes as training set, 10% as validation set, and 10% as test set. Hence, part of the genes of *E. coli* are used for training.
- **Gustafson et al.**: Gustafson et al. [85] use sequence-derived and experimental features to classify essential genes in *E. coli* and *S. cerevisiae*. They separately train a naïve Bayes classifier for each species, using half of its essential genes and half of its non-essential genes as training set. For *E. coli*, they only use features derived from the sequence. The AU-ROC is computed on all genes, irrespective of whether they were used for training.
- **Evo 1**: Evo 1 [24] is a genomic foundation model that is trained in a self-supervised way on nucleotide sequences. Evo 1 has 7 billion parameters and a context length of up to 131,072 tokens. It was trained on prokaryotic sequences and was shown to be able to predict gene essentiality in a zero-shot fashion. Specifically, the model is given as input a 8192-nucleotide long context window centered in a protein-coding gene. Then a perturbation consisting of several stop codons (TAATAATAATAGTGA) is inserted at 12 bp after the beginning of the gene. The two sequences are passed through the model, which outputs a likelihood for each sequence. The ratio of the likelihoods is then used to predict essential genes. Nguyen et al. [24] used the model to make predictions for 56 bacterial species, including *E. coli*, and 2 phage species. The AU-ROC shown in Figure S4 was computed using the version of Evo 1 with 131k nucleotide context.
- **Evo 2**: Evo 2 [25] is a family of foundation models, with 7 billion or 40 billion parameters, trained at different context window lengths. It is trained using both prokaryotic and eukaryotic species. The gene essentiality prediction pipeline used for Evo 2 is identical to the one used for Evo 1. They report predictions only for prokaryotic species and for human lncRNAs (for which they use a different genetic perturbation, scrambling 100 bp portions of DNA at specific positions given by Cas13 guide sequence binding sites). The AU-ROC shown in Figure S4 was computed using the version of Evo 2 with 7B parameters and 8k nucleotide context.

Figure S4 compares ProteomeLM-Ess to the following methods on *S. cerevisiae*:

- **Yue et al.**: Yue et al. [86] use a deep learning method that combines different sources of information to predict gene essentiality. They separately construct embedding vectors for gene expression, PPI network, and subcellular localization. They then concatenate the three embedding vectors and use them as input for a one-layer neural network classifier with sigmoid activation. The model is used only on *S. cerevisiae* data, with 60% of the data used as training set, 20% as validation set and 20% for testing.
- **Gustafson et al.**: see above for method description. For *S. cerevisiae*, they both train a classifier using only sequence-derived features *(sequence)* and another one where these are combined with features obtained from experimental data *(all)*. Importantly, as for *E. coli*, some of the genes of *S. cerevisiae* are used for training and others for testing, and the AU-ROC shown in Figure S4 is computed on all genes, irrespective of whether they were used for training.
- **Saha&Heber**: Saha and Heber [87] use 1098 essential and 1098 non-essential genes of *S. cerevisiae* as training data for their classifier. They predict gene essentiality by combining a *k* nearest neighbor and a support vector machine classifier, which take as input features derived from the sequence, from protein-protein interaction data, as well as from the comparison between yeast data and data from other species.
- **OGN**: Zhang et al. [88] propose a mathematical model that combines measures of correlation in expression data with PPI network features. Their model outputs a parameter that is used to rank genes from most likely essential to least likely essential. They test their model on *S. cerevisiae* data.
- **Evo 1**: see above for method description. Recall that the model was trained and evaluated only on data from prokaryotic species. Hence, we performed our own evaluation of gene essentiality in *S. cerevisiae*, using the same perturbation as for prokaryotes (see above). Note however that this is an out-of-distribution task for this model.
- **Evo 2**: see above for a description of the method. Recall that, contrary to Evo 1, the model’s training set includes eukaryotes. However, gene essentiality prediction is not reported in Ref. [25] for these species. Hence, as for Evo 1, we performed our own evaluation of gene essentiality in *S. cerevisiae*. Note however that the choice of perturbation might not be optimal for eukaryotes.

### 6 Comparison of continuous and discrete functional encodings

Could the functional encoding used by ProteomeLM, which averages the ESM-C embeddings of proteins within each orthologous group, be replaced by a simpler discrete representation of orthologous group identity? To address this question, we trained an alternative version of ProteomeLM, which we refer to as ProteomeLM-Discrete, in which the continuous functional encoding is replaced by a learnable table indexed by OrthoDB group identifiers. Concretely, each orthologous group in the OrthoDB database [67] is assigned a unique integer index, and the model learns a fixed-dimensional embedding vector for each index from scratch during training. This design is analogous to the positional encoding usually employed in language models [114], where each position (here, each orthologous group identity) is represented by a learned vector. The rest of the architecture, including the transformer encoder, the masked language modeling objective, and the prediction heads, is kept identical to the standard ProteomeLM, in order to ensure a fair comparison.

#### Architecture details

We constructed an OrthoDB vocabulary comprising all orthologous group identifiers present in the training set, yielding a vocabulary of approximately 1.7 million entries. Each entry is mapped to a learnable embedding of dimension 1152 (matching the ESM-C embedding dimension), initialized with a normal distribution. An additional index is reserved for proteins with no known orthologous group assignment (`<UNK>`). For each protein, the orthologous group at the broadest taxonomic level is selected. ProteomeLM-Discrete was trained for 210 epochs under identical hardware and hyperparameter conditions as ProteomeLM-S (36M parameters), using the MSE loss, since the motivation for the polar loss (preventing collapse toward the functional encoding, see Methods and Section 7) does not apply when the functional encoding is a learned discrete vector unrelated to the target ESM-C embedding.

#### Results

We evaluate unsupervised PPI recovery by ProteomeLM-Discrete and compare it to the standard ProteomeLM-S on the benchmark distinguishing direct interactions, same-complex interactions and coexpression associations described in Section 1. For each model, we compute the AUROC of each individual attention head at distinguishing positive pairs from random pairs, across three species: *E. coli*, *S. cerevisiae*, and *H. sapiens*.

Figure S5 shows the per-head AUROC values, sorted in descending order, for both models across the three interaction types. ProteomeLM-Discrete (dashed lines) captures PPI signal across all species and interaction types, confirming that orthologous group identity alone provides a useful signal for learning inter-protein dependencies. However, our standard ProteomeLM with continuous functional encoding (solid lines) consistently outperforms ProteomeLM-Discrete. The gap is more pronounced in *E. coli*, where the best heads of ProteomeLM reach an AUROC of 0.87 for direct interactions, compared to 0.70 for ProteomeLM-Discrete. In *S. cerevisiae* and *H. sapiens*, ProteomeLM also possesses an advantage over ProteomeLM-Discrete, though the gap is smaller for coexpression associations than for direct or same-complex interactions.

**Figure S5:**
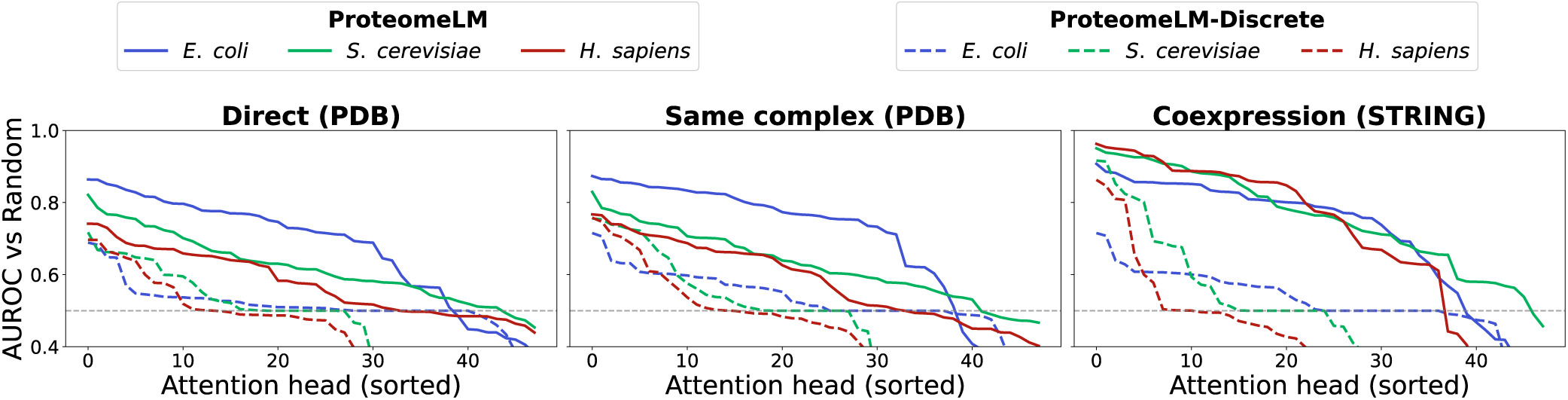
Comparison of continuous and discrete functional encodings for unsupervised PPI prediction. Per-head AUROC for distinguishing interacting from random protein pairs, sorted in descending order, for the standard ProteomeLM-S with continuous functional encoding and for ProteomeLM-Discrete with learnable OrthoDB group embeddings. Results are shown for three interaction types: direct interactions (PDB), same-complex interactions (PDB), and coexpression (STRING), see Section 1, across three species: *E. coli*, *S. cerevisiae*, and *H. sapiens*. Both models comprise 48 attention heads (6 layers × 8 heads).

#### Interpretation

These results indicate that our continuous functional encoding is beneficial beyond simply providing orthologous group identity. By encoding the average ESM-C embedding of each group, the functional encoding injects protein family-level sequence information into the model’s input, which can be used by the model to better resolve inter-protein dependencies. The discrete encoding, while sufficient to capture broad co-occurrence patterns (as reflected in its reasonable coexpression recovery), lacks the fine-grained biochemical signal carried by ESM-C embeddings, which appears important in particular for distinguishing physical interactions. In addition, with a discrete encoding, the model cannot as efficiently transfer signal from one family to the other, which is critical when working with small families and with protein sequences with no assigned OrthoDB group. These results validate our design choice of using continuous, ESM-C-derived functional encodings.

### 7 Theoretical motivation of the polar loss used to train ProteomeLM

Let *x* denote the true protein embedding and *x̅* the functional encoding. We define the true residual vector as *r* = *x* − *x̅*. Similarly, let *x̂* be the predicted embedding and *r̂* = *x̂* − *x̅* be the predicted residual.

#### Gradient collapse with the mean-squared-error loss

Let us write the standard mean-squared-error (MSE) loss between *x̂* and *x* as a function of the residuals:

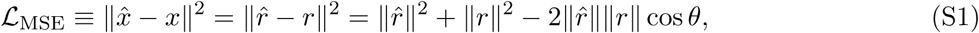

where ∥.∥ denotes the Euclidean norm, and *θ* is the angle between *r̂* and *r*. The gradient of this loss with respect to the angular alignment *θ* between *r̂* and *r* is given by:

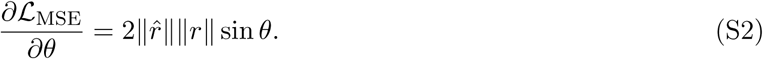

Eq. S2 reveals that the signal for learning the correct direction is scaled by the product of the magnitudes of the true residual ∥*r*∥ and of the predicted residual ∥*r̂*∥. In our setting, *x* and *x̅* are highly correlated, meaning that the true residual magnitude ∥*r*∥ is small. Moreover, if the model defaults to the functional encoding for the protein embedding (i.e. *x̂* → *x̅*), then ∥*r̂*∥ → 0. Consequently, the angular gradient in Eq. S2 vanishes. This prevents the model from learning the specific deviations of *x* from *x̅*, and leads to the degenerate solution described in the main text.

#### Derivation of the polar loss via maximum likelihood estimation under magnitude-direction decoupling

In the maximum likelihood spirit, let us consider the true residual *r* as a random variable conditioned on the model prediction *r̂*. To address the challenge identified above, let us further decouple the magnitude and the direction of the residual. To this end, we assume independent distributions for its radial (magnitude) and angular (directional) components in polar coordinates:

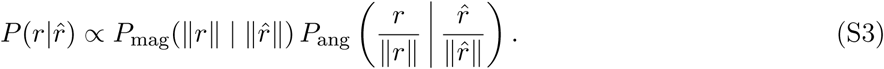

Let us now make simple assumptions on the explicit forms of the two distributions *P*_mag_ and *P*_ang_.

*Magnitude Component:* We assume that the magnitude ∥*r*∥ follows a normal distribution (denoted by N) centered at the predicted magnitude ∥*r̂*∥ with variance *σ*^2^:

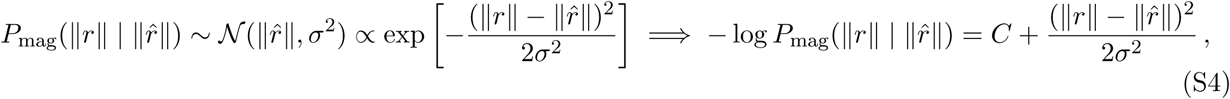

where *C* is a constant.

*Directional Component:* We further assume that the unit direction vector *u* = *r/*∥*r*∥ follows a von Mises-Fisher distribution (denoted by vMF) centered at the predicted direction *û* = *r̂*/∥*r̂*∥ and with concentration parameter *κ*:

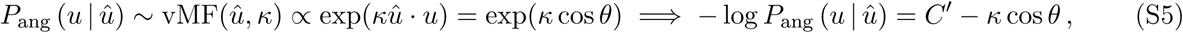

where *û* · *u* denotes the canonical scalar product of *û* and *u*, and *C^′^* is a constant.

Under the two assumptions above, the negative log-likelihood of *P* (*r*|*r̂*) reads:

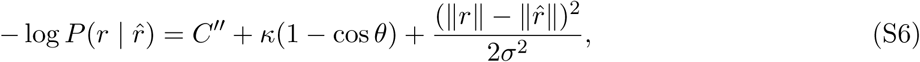

where *C^′′^* is a constant. Thus, under our assumptions, maximizing the likelihood of the data *r* under the model *r̂* amounts to minimizing a loss of the following form:

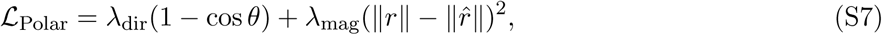

where *λ*_dir_ and *λ*_mag_ are hyperparameters corresponding to the inverse variances of the directional and magnitude noise, respectively. In practice, we use *λ*_dir_ = *λ*_mag_ = 1 in our polar loss, as a pragmatic choice. Crucially, the angular gradient of this loss is proportional to sin *θ* but *independent* of ∥*r̂*∥. This decoupling prevents gradients for directional alignment from vanishing when the magnitude is small, thus avoiding the mode collapse identified above. Furthermore, our polar loss is minimized if and only if *x̂* = *x*. Empirical validation of the polar loss against alternatives is shown in Figure 6D.

## 8 Supplementary figures

**Figure S6:**
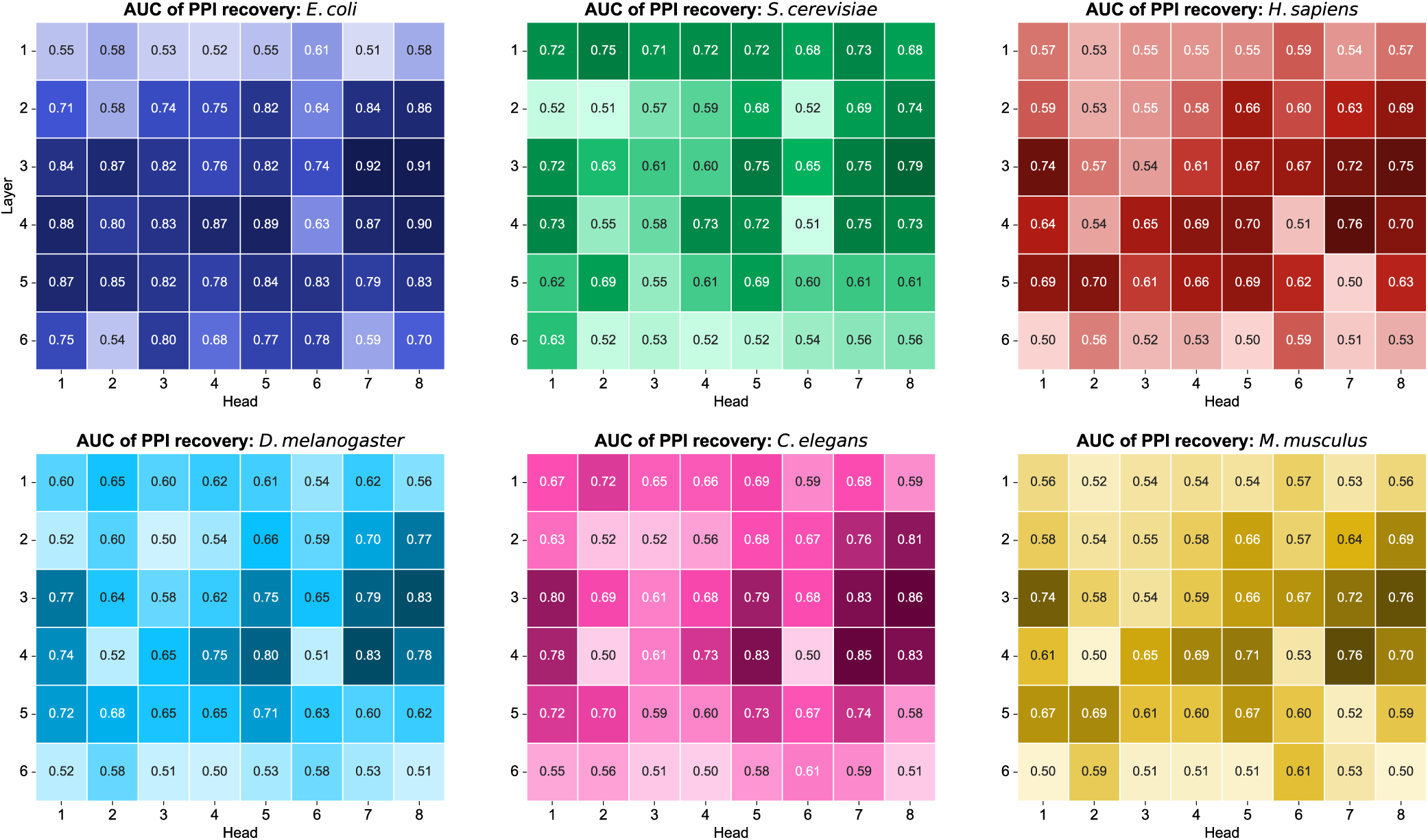
Attention head-wise PPI recovery across species in ProteomeLM-S. The area under the ROC curve (AUC) for protein-protein interaction (PPI) prediction using each individual attention head of ProteomeLM-S (6layers, 8 heads) is reported across six species: *E*. *coli*, *S. cerevisiae*, *H. sapiens*, *D. melanogaster*, *C. elegans*, and *M. musculus*. Each heatmap shows the AUC for one species, with values computed separately for each attention head and layer. Attention heads from intermediate layers (particularly layer 3) consistently exhibit high predictive power, especially in *E. coli*. The first three panels (top) are also shown in Figure 2, and are reproduced here to facilitate inter-species comparison.

**Figure S7:**
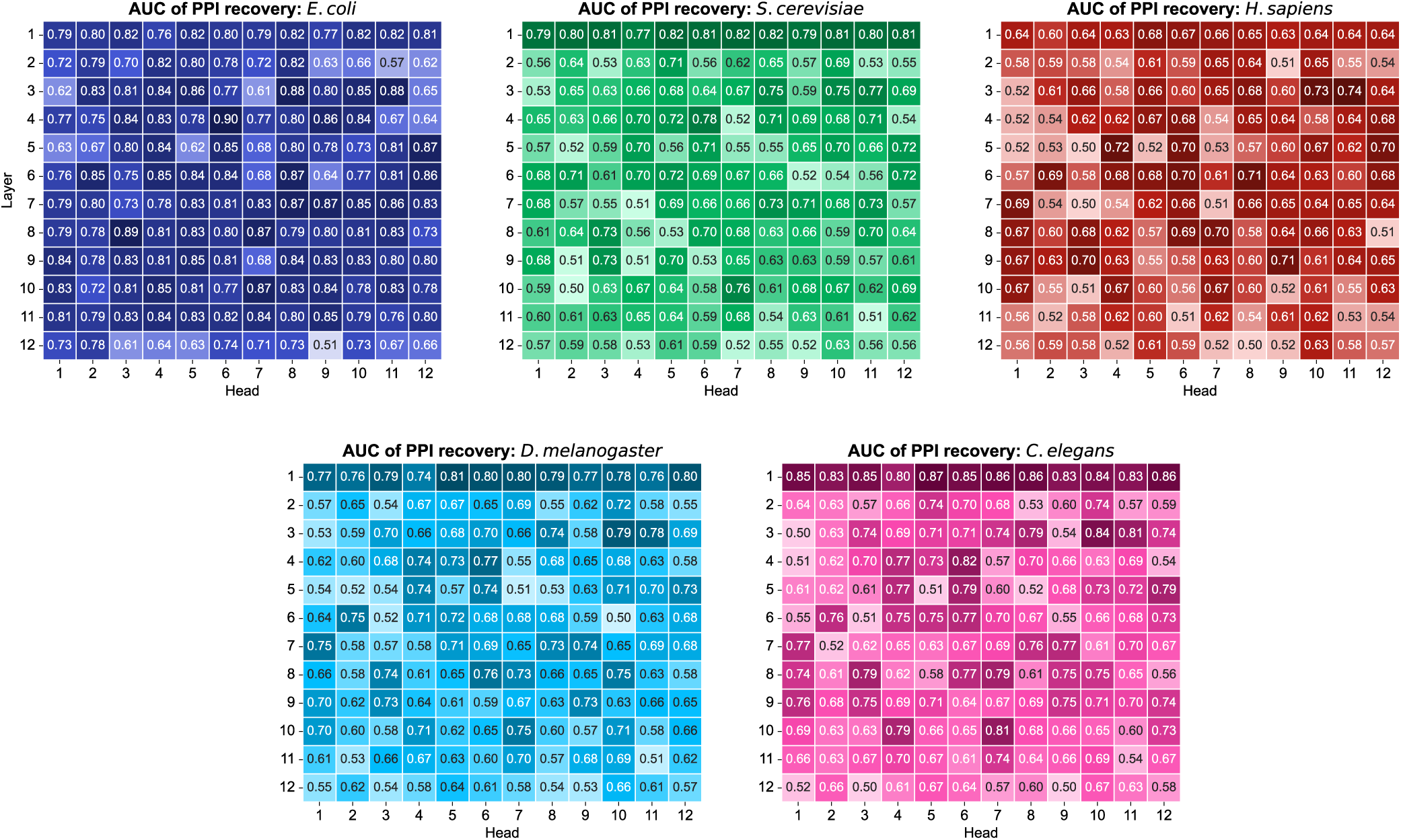
Attention head-wise AUC of PPI recovery across species in ProteomeLM-M. The AUC of protein-protein interaction (PPI) prediction is reported for each of the 12attention head from all 12 layers of ProteomeLM-M (112M parameters), for five species: *E*. *coli*, *S. cerevisiae*, *H. sapiens*, *D. melanogaster*, and *C. elegans*. As in ProteomeLM-S (Figure S6), central layers tend to concentrate the most predictive heads. However, we note that the first layer is also a good predictor of PPI, in particular in eukaryotes. Note that the proteome of *M. musculus* is too long to allow the use of ProteomeLM-M.

**Figure S8:**
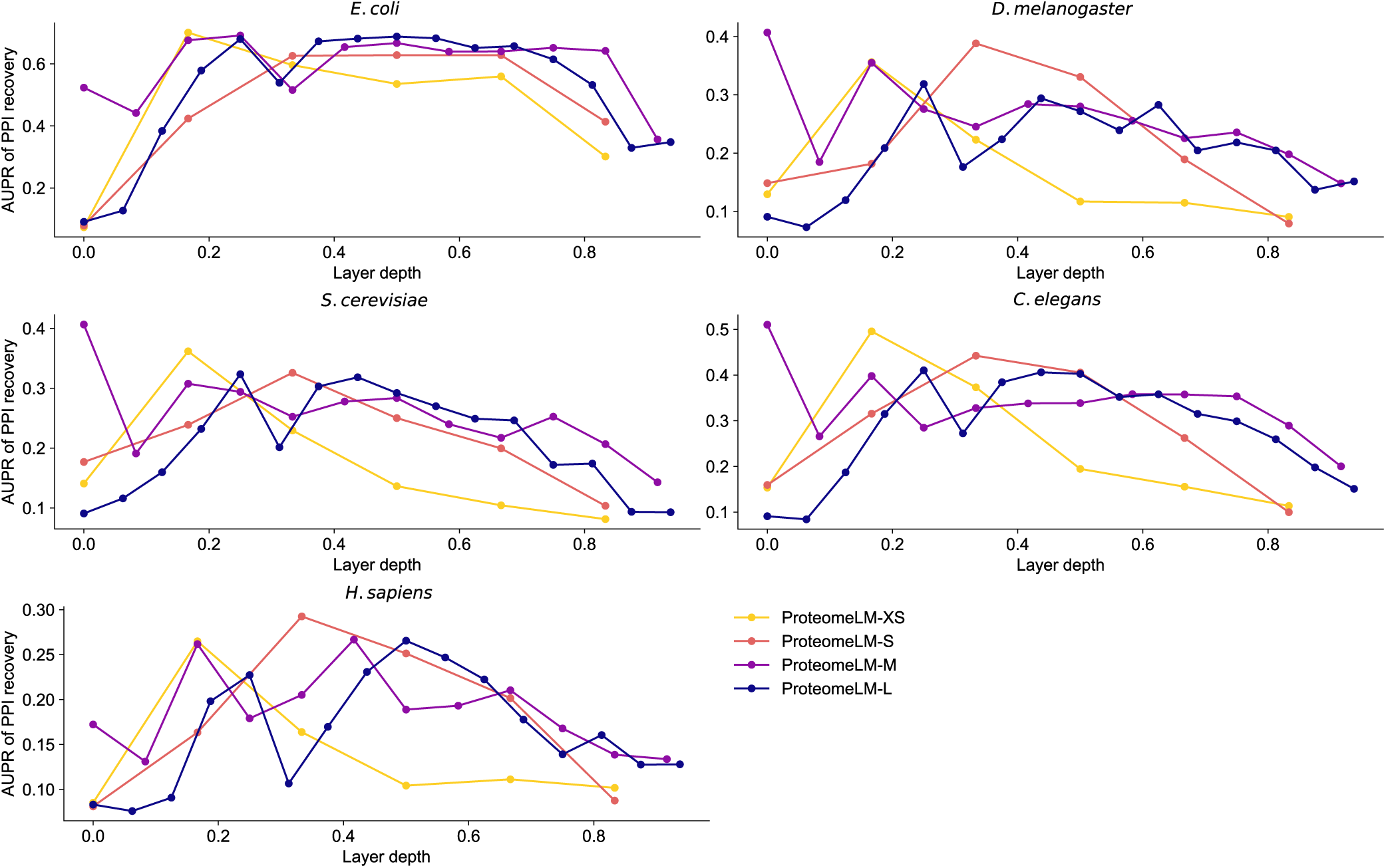
Unsupervised PPI prediction by ProteomeLM across model sizes and layers. The area under the precision-recall curve (AUPR) for unsupervised PPI prediction using summed attention scores is shown for each layer, across four model sizes (XS, S, M, L) and five species: *E. coli*, *S. cerevisiae*, *H. sapiens*, *D. melanogaster*, and *C. elegans*. To facilitate comparison between models with different sizes, the AUPR is plotted versus the normalized layer depth. Performance peaks at intermediate layers across species and model sizes.

**Figure S9:**
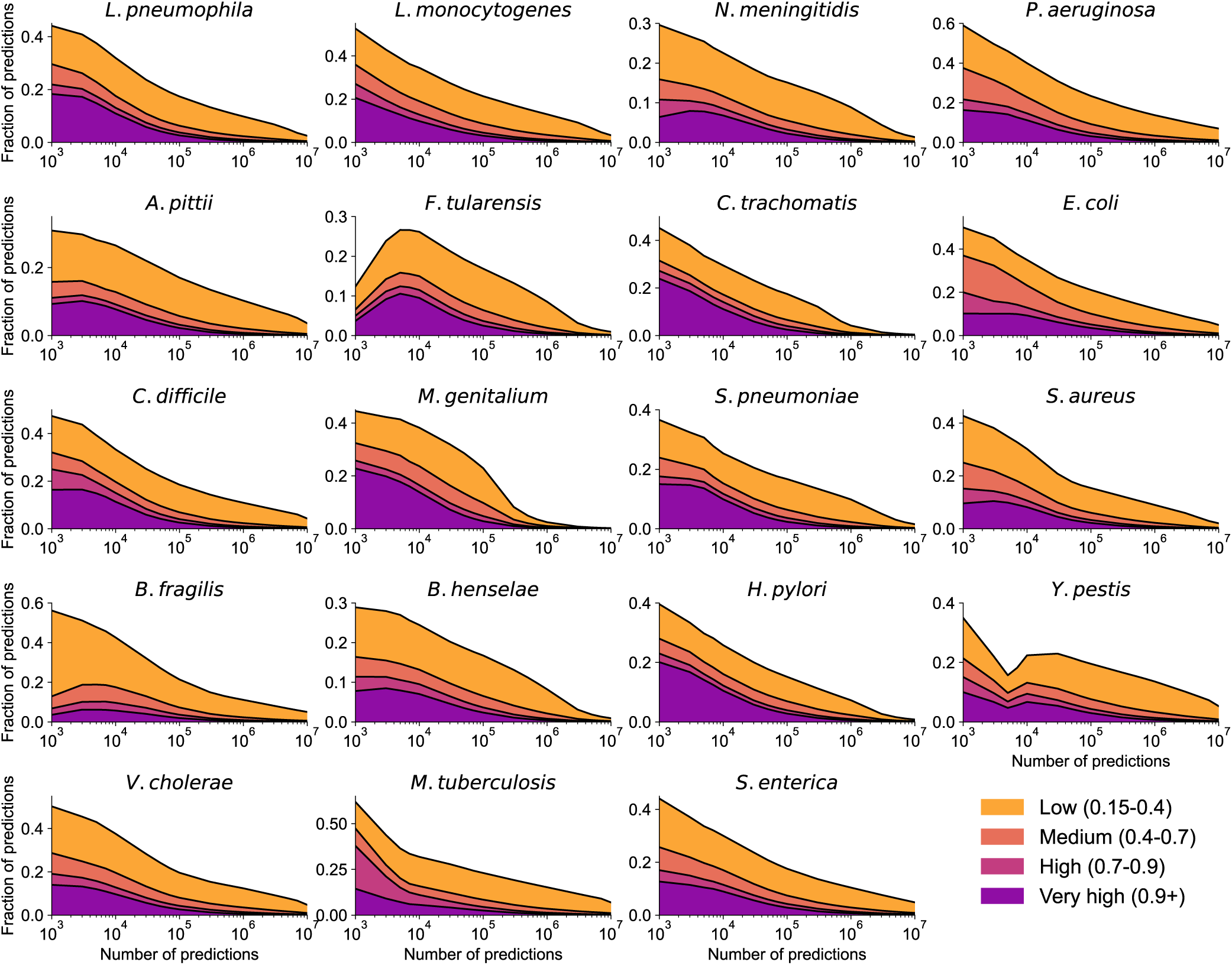
Accuracy of fast PPI screening by ProteomeLM across 19 pathogens. The fraction of top-scoring predicted PPI that correspond to known interactions in the STRING database is shown across 19 human bacterial pathogens. Predictions are sorted by the ProteomeLM classifier score, and their cumulative fraction in each of four bins of STRING confidence score is plotted as a function of the number of top-ranked predictions considered. Across most pathogens, a substantial fraction of high-scoring predictions correspond to known or high-confidence interactions. The 19 pathogens considered are the same as in Figure 3, and aggregated results are shown in Figure 3D.

**Figure S10:**
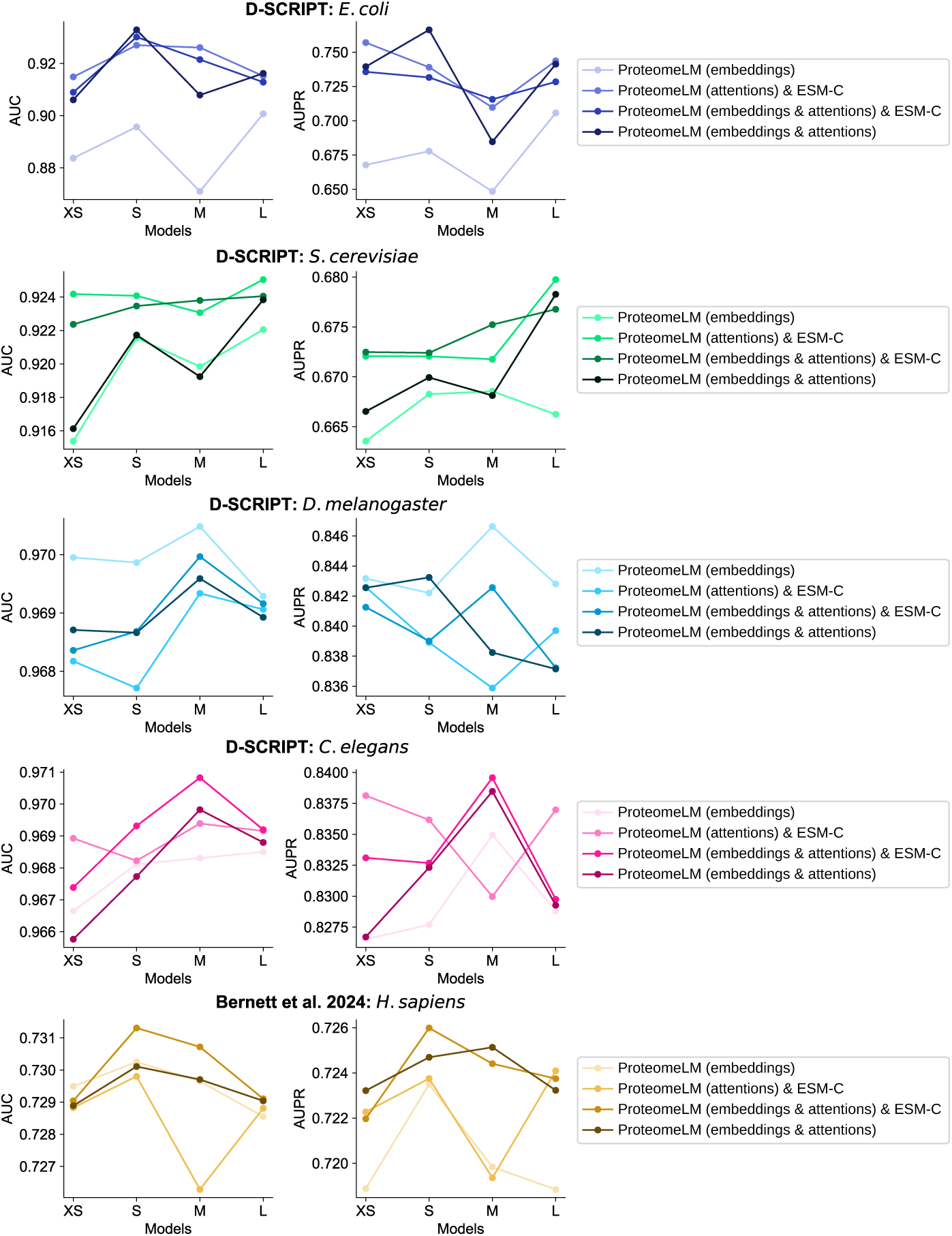
Impact of embeddings and attention values on supervised PPI prediction. Evaluation of supervised PPI prediction across four ProteomeLM model sizes (XS, S, M, L), on five benchmarks: the D-SCRIPT datasets for *E. coli*, *S. cerevisiae*, *D. melanogaster*, *C. elegans*, and the dataset from Ref. [76] for *H. sapiens*. For each of these benchmarks, both the AUC (left) and the AUPR (right) are reported using four different input feature configurations for PPI prediction: ProteomeLM embeddings only; ESM-C embeddings and ProteomeLM attention; ESM-C embeddings, ProteomeLM embeddings and ProteomeLM attention; and finally, ProteomeLM embeddings and ProteomeLM attention. The latter is the configuration we retained throughout, and called ProteomeLM-PPI. Indeed, we observe here that combining ProteomeLM embeddings and ProteomeLM attention yields the best or near-best performance across species and metrics.

**Figure S11:**
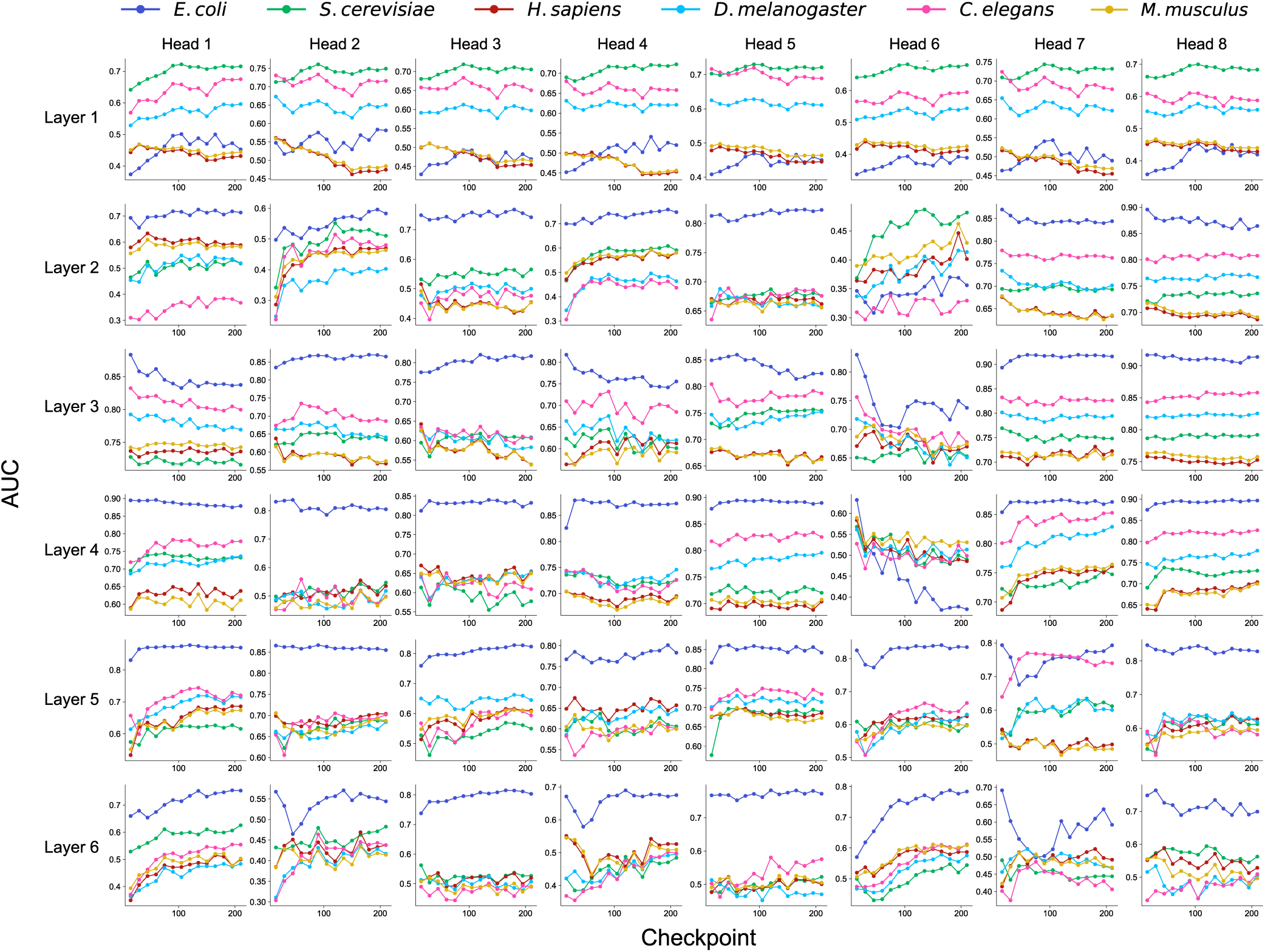
Unsupervised recovery of PPI by ProtomeLM-S attention heads during training. The area under the ROC curve (AUC) for unsupervised PPI recovery is shown versus training checkpoints for each attention head of ProteomeLM-S. Curves are shown separately for six species: *E. coli*, *S. cerevisiae*, *H. sapiens*, *D. melanogaster*, *C. elegans*, and *M. musculus*. Different heads exhibit different dynamics and specialization. Some become increasingly precise on *E. coli* (e.g. layer 6, head 3), while others (e.g. layer 1, head 1) are more specialized on eukaryotes. Many heads exhibit increasingly good PPI recovery along training.

## Notes

https://github.com/Bitbol-Lab/ProteomeLM

### Competing Interest Statement

The authors have declared no competing interest.

### Summary of Updates

Diverse revision and new results were added following reviewers recommendation after a first round of reviews: - We have added results on distinguishing direct PPIs from functional links - We have added examples on resolved structural complexes - We have added more benchmarking on gene essentiality - We have improved some explanations in the methods

## References

[1] E. C. Alley, G. Khimulya, S. Biswas, M. AlQuraishi, and G. M. Church. Unified rational protein engineering with sequence-based deep representation learning. Nature Methods, 16(12):1315–1322, 2019.

[2] Ahmed Elnaggar, Michael Heinzinger, Christian Dallago, Ghalia Rehawi, Yu Wang, Llion Jones, Tom Gibbs, Tamas Feher, Christoph Angerer, Martin Steinegger, Debsindhu Bhowmik, and Burkhard Rost. ProtTrans: Towards cracking the language of life’s code through self-supervised deep learning and high performance computing. IEEE Transactions on Pattern Analysis and Machine Intelligence, 44(10):7112–7127, 2022.

[3] Alexander Rives, Joshua Meier, Tom Sercu, Siddharth Goyal, Zeming Lin, Jason Liu, Demi Guo, Myle Ott, C. Lawrence Zitnick, Jerry Ma, and Rob Fergus. Biological structure and function emerge from scaling unsupervised learning to 250 million protein sequences. Proc. Natl. Acad. Sci. USA, 118(15):e2016239118, 2021.

[4] Roshan Rao, Joshua Meier, Tom Sercu, Sergey Ovchinnikov, and Alexander Rives. Transformer protein language models are unsupervised structure learners. In International Conference on Learning Representations, 2021.

[5] Jesse Vig, Ali Madani, Lav R. Varshney, Caiming Xiong, Richard Socher, and Nazneen Rajani. BERTology meets biology: Interpreting attention in protein language models. In International Conference on Learning Representations, 2021.

[6] Ali Madani, Bryan McCann, Nikhil Naik, Nitish Shirish Keskar, Namrata Anand, Raphael R. Eguchi, Po-Ssu Huang, and Richard Socher. ProGen: Language modeling for protein generation. bioRxiv, page 2020.03.07.982272, 2020.

[7] A. Madani, B. Krause, E. R. Greene, S. Subramanian, B. P. Mohr, J. M. Holton, J. L. Olmos, C. Xiong, Z. Z. Sun, R. Socher, J. S. Fraser, and N. Naik. Large language models generate functional protein sequences across diverse families. Nat. Biotechnol., 41(8):1099–1106, 2023.

[8] Noelia Ferruz, Steffen Schmidt, and Birte Höcker. ProtGPT2 is a deep unsupervised language model for protein design. Nature Communications, 13(1):4348, 2022.

[9] Roshan M Rao, Jason Liu, Robert Verkuil, Joshua Meier, John Canny, Pieter Abbeel, Tom Sercu, and Alexander Rives. MSA Transformer. Proceedings of the 38th International Conference on Machine Learning, 139:8844–8856, 2021.

[10] Zeming Lin, Halil Akin, Roshan Rao, Brian Hie, Zhongkai Zhu, Wenting Lu, Nikita Smetanin, Robert Verkuil, Ori Kabeli, Yaniv Shmueli, Allan Dos Santos Costa, Maryam Fazel-Zarandi, Tom Sercu, Salvatore Candido, and Alexander Rives. Evolutionary-scale prediction of atomic-level protein structure with a language model. Science, 379(6637):1123–1130, 2023.

[11] Nadav Brandes, Dan Ofer, Yam Peleg, Nadav Rappoport, and Michal Linial. ProteinBERT: a universal deep-learning model of protein sequence and function. Bioinformatics, 38(8):2102–2110, 2022.

[12] ESM Team et al. ESM Cambrian: Revealing the mysteries of proteins with unsupervised learning. Evolutionary Scale Website, https://www.evolutionaryscale.ai/blog/esm-cambrian, 2024.

[13] Vineet Thumuluri, José Juan Almagro Armenteros, Alexander Rosenberg Johansen, Henrik Nielsen, and Ole Winther. DeepLoc 2.0: multi-label subcellular localization prediction using protein language models. Nucleic acids research, 50(W1):W228–W234, 2022.

[14] Joshua Meier, Roshan Rao, Robert Verkuil, Jason Liu, Tom Sercu, and Alexander Rives. Language models enable zero-shot prediction of the effects of mutations on protein function. NeurIPS, 2021.

[15] Pranav Kantroo, Günter P. Wagner, and Benjamin B. Machta. Pseudo-perplexity in one fell swoop for protein fitness estimation. arXiv, page 2407.07265, 2024.

[16] D. Miller, A. Stern, and D. Burstein. Deciphering microbial gene function using natural language processing. Nat. Commun., 13(1):5731, 2022.

[17] Yanrong Ji, Zhihan Zhou, Han Liu, and Ramana V Davuluri. DNABERT: pre-trained Bidirectional Encoder Representations from Transformers model for DNA-language in genome. Bioinformatics, 37(15):2112–2120, 2021.

[18] Hugo Dalla-Torre, Liam Gonzalez, Javier Mendoza-Revilla, Nicolas Lopez Carranza, Adam Henryk Grzywaczewski, Francesco Oteri, Christian Dallago, Evan Trop, Bernardo P. de Almeida, Hassan Sirelkhatim, Guillaume Richard, Marcin Skwark, Karim Beguir, Marie Lopez, and Thomas Pierrot. Nucleotide Transformer: building and evaluating robust foundation models for human genomics. Nature Methods, 22(2):287–297, 2025.

[19] Eric Nguyen, Michael Poli, Marjan Faizi, Armin Thomas, Michael Wornow, Callum Birch-Sykes, Stefano Massaroli, Aman Patel, Clayton Rabideau, Yoshua Bengio, Stefano Ermon, Stephen A. Baccus, and Chris Ré. HyenaDNA: Long-range genomic sequence modeling at single nucleotide resolution. Advances in neural information processing systems, 36:43177–43201, 2023.

[20] Gonzalo Benegas, Carlos Albors, Alan J. Aw, Chengzhong Ye, and Yun S. Song. GPN-MSA: an alignment-based DNA language model for genome-wide variant effect prediction. bioRxiv, page 2023.10.10.561776, 2024.

[21] Yunha Hwang, Andre L. Cornman, Elizabeth H. Kellogg, Sergey Ovchinnikov, and Peter R. Girguis. Genomic language model predicts protein co-regulation and function. Nature Communications, 15(1):2880, 2024.

[22] Andre Cornman, Jacob West-Roberts, Antonio Pedro Camargo, Simon Roux, Martin Beracochea, Milot Mirdita, Sergey Ovchinnikov, and Yunha Hwang. The OMG dataset: An Open MetaGenomic corpus for mixed-modality genomic language modeling. bioRxiv, page 2024.08.14.607850, 2024.

[23] Bin Shao and Jiawei Yan. A long-context language model for deciphering and generating bacteriophage genomes. Nature Communications, 15(1):9392, 2024.

[24] Eric Nguyen, Michael Poli, Matthew G. Durrant, Brian Kang, Dhruva Katrekar, David B. Li, Liam J. Bartie, Armin W. Thomas, Samuel H. King, Garyk Brixi, Jeremy Sullivan, Madelena Y. Ng, Ashley Lewis, Aaron Lou, Stefano Ermon, Stephen A. Baccus, Tina Hernandez-Boussard, Christopher Ré, Patrick D. Hsu, and Brian L. Hie. Sequence modeling and design from molecular to genome scale with Evo. Science, 386(6723):eado9336, 2024.

[25] Garyk Brixi, Matthew G. Durrant, Jerome Ku, Michael Poli, Greg Brockman, Daniel Chang, Gabriel A. Gonzalez, Samuel H. King, David B. Li, Aditi T. Merchant, Mohsen Naghipourfar, Eric Nguyen, Chiara Ricci-Tam, David W. Romero, Gwanggyu Sun, Ali Taghibakshi, Anton Vorontsov, Brandon Yang, Myra Deng, Liv Gorton, Nam Nguyen, Nicholas K. Wang, Etowah Adams, Stephen A. Baccus, Steven Dillmann, Stefano Ermon, Daniel Guo, Rajesh Ilango, Ken Janik, Amy X. Lu, Reshma Mehta, Mohammad R. K. Mofrad, Madelena Y. Ng, Jaspreet Pannu, Christopher Ré, Jonathan C. Schmok, John St John, Jeremy Sullivan, Kevin Zhu, Greg Zynda, Daniel Balsam, Patrick Collison, Anthony B. Costa, Tina Hernandez-Boussard, Eric Ho, Ming-Yu Liu, Thomas McGrath, Kimberly Powell, Dave P. Burke, Hani Goodarzi, Patrick D. Hsu, and Brian L. Hie. Genome modeling and design across all domains of life with Evo 2. bioRxiv, page 2025.02.18.638918, 2025.

[26] Žiga Avsec, Natasha Latysheva, Jun Cheng, Guido Novati, Kyle R. Taylor, Tom Ward, Clare Bycroft, Lauren Nicolaisen, Eirini Arvaniti, Joshua Pan, Raina Thomas, Vincent Dutordoir, Matteo Perino, Soham De, Alexander Karollus, Adam Gayoso, Toby Sargeant, Anne Mottram, Lai Hong Wong, Pavol Drotár, Adam Kosiorek, Andrew Senior, Richard Tanburn, Taylor Applebaum, Souradeep Basu, Demis Hassabis, and Pushmeet Kohli. AlphaGenome: advancing regulatory variant effect prediction with a unified DNA sequence model. bioRxiv, page 2025.06.25.661532, 2025.

[27] Samuel Sledzieski, Rohit Singh, Lenore Cowen, and Bonnie Berger. D-SCRIPT translates genome to phenome with sequence-based, structure-aware, genome-scale predictions of protein-protein interactions. Cell Systems, 12(10):969–982, 2021.

[28] S. V. Rajagopala, P. Sikorski, A. Kumar, R. Mosca, J. Vlasblom, R. Arnold, J. Franca-Koh, S. B. Pakala, S. Phanse, A. Ceol, R. Hauser, G. Siszler, S. Wuchty, A. Emili, M. Babu, P. Aloy, R. Pieper, and P. Uetz. The binary protein-protein interaction landscape of Escherichia coli. Nat. Biotechnol., 32(3):285–290, 2014.

[29] Rose Oughtred, Jennifer Rust, Christie Chang, Bobby-Joe Breitkreutz, Chris Stark, Andrew Willems, Lorrie Boucher, Genie Leung, Nadine Kolas, Frederick Zhang, Sonam Dolma, Jasmin Coulombe-Huntington, Andrew Chatr-Aryamontri, Kara Dolinski, and Mike Tyers. The BioGRID database: A comprehensive biomedical resource of curated protein, genetic, and chemical interactions. Protein Science, 30(1):187–200, 2021.

[30] Peter D Karp, Wai Kit Ong, Suzanne Paley, Richard Billington, Ron Caspi, Carol Fulcher, Anamika Kothari, Markus Krummenacker, Mario Latendresse, Peter E Midford, Pallavi Subhraveti, Socorro Gama-Castro, Luis Muñiz-Rascado, César Bonavides-Martinez, Alberto Santos-Zavaleta, Amanda Mackie, Julio Collado-Vides, Ingrid M. Keseler, and Ian Paulsen. The EcoCyc database. EcoSal Plus, 8(1):10–1128, 2018.

[31] Noemi Del Toro, Anjali Shrivastava, Eliot Ragueneau, Birgit Meldal, Colin Combe, Elisabet Barrera, Livia Perfetto, Karyn How, Prashansa Ratan, Gautam Shirodkar, et al. The IntAct database: efficient access to fine-grained molecular interaction data. Nucleic acids research, 50(D1):D648–D653, 2022.

[32] Rodrigo V. Honorato, Mikael E. Trellet, Brian Jiménez-García, Jörg J. Schaarschmidt, Marco Giulini, Victor Reys, Panagiotis I. Koukos, João P. G. L. M. Rodrigues, Ezgi Karaca, Gydo C. P. van Zundert, Jorge Roel-Touris, Charlotte W. van Noort, Zuzana Jandová, Adrien S. J. Melquiond, and Alexandre M. J. J. Bonvin. The HADDOCK2.4 web server for integrative modeling of biomolecular complexes. Nature Protocols, 19(11):3219–3241, 2024.

[33] Mohamed Amine Ketata, Cedrik Laue, Ruslan Mammadov, Hannes Stark, Menghua Wu, Gabriele Corso, Céline Marquet, Regina Barzilay, and Tommi S Jaakkola. DiffDock-PP: Rigid protein-protein docking with diffusion models. In ICLR 2023-Machine Learning for Drug Discovery workshop, 2023.

[34] Azam Shirali, Vitalii Stebliankin, Ukesh Karki, Jimeng Shi, Prem Chapagain, and Giri Narasimhan. A comprehensive survey of scoring functions for protein docking models. BMC bioinformatics, 26(1):25, 2025.

[35] Ameya Harmalkar, Sergey Lyskov, and Jeffrey J Gray. Reliable protein–protein docking with AlphaFold, Rosetta, and replica exchange. Elife, 13:RP94029, 2025.

[36] Richard Evans, Michael O’Neill, Alexander Pritzel, Natasha Antropova, Andrew Senior, Tim Green, Augustin Žídek, Russ Bates, Sam Blackwell, Jason Yim, Olaf Ronneberger, Sebastian Bodenstein, Michal Zielinski, Alex Bridgland, Anna Potapenko, Andrew Cowie, Kathryn Tunyasuvunakool, Rishub Jain, Ellen Clancy, Pushmeet Kohli, John Jumper, and Demis Hassabis. Protein complex prediction with AlphaFold-Multimer. bioRxiv, page 2021.10.04.463034, 2021.

[37] Patrick Bryant, Gabriele Pozzati, and Arne Elofsson. Improved prediction of protein-protein interactions using AlphaFold2. Nature communications, 13(1):1265, 2022.

[38] U. Göbel, C. Sander, R. Schneider, and A. Valencia. Correlated mutations and residue contacts in proteins. Proteins, 18(4):309–317, 1994.

[39] G. Casari, C. Sander, and A. Valencia. A method to predict functional residues in proteins. Nat. Struct. Biol., 2(2):171–178, 1995.

[40] Michael Socolich, Steve W. Lockless, William P. Russ, Heather Lee, Kevin H. Gardner, and Rama Ranganathan. Evolutionary information for specifying a protein fold. Nature, 437(7058):512–518, 2005.

[41] S. D. Dunn, L. M. Wahl, and G. B. Gloor. Mutual information without the influence of phylogeny or entropy dramatically improves residue contact prediction. Bioinformatics, 24(3):333–340, 2008.

[42] M. Weigt, R. A. White, H. Szurmant, J. A. Hoch, and T. Hwa. Identification of direct residue contacts in protein-protein interaction by message passing. Proc. Natl. Acad. Sci. USA, 106(1):67–72, 2009.

[43] D. S. Marks, L. J. Colwell, R. Sheridan, T. A. Hopf, A. Pagnani, R. Zecchina, and C. Sander. Protein 3D structure computed from evolutionary sequence variation. PLOS ONE, 6(12):e28766, 2011.

[44] F. Morcos, A. Pagnani, B. Lunt, A. Bertolino, D. S. Marks, C. Sander, R. Zecchina, J. N. Onuchic, T. Hwa, and M. Weigt. Direct-coupling analysis of residue coevolution captures native contacts across many protein families. Proc. Natl. Acad. Sci. USA, 108(49):E1293–1301, 2011.

[45] Anne-Florence Bitbol, Robert S Dwyer, Lucy J Colwell, and Ned S Wingreen. Inferring interaction partners from protein sequences. Proc. Natl. Acad. Sci. USA, 113(43):12180–12185, 2016.

[46] T. Gueudre, C. Baldassi, M. Zamparo, M. Weigt, and A. Pagnani. Simultaneous identification of specifically interacting paralogs and interprotein contacts by direct coupling analysis. Proc. Natl. Acad. Sci. USA, 113(43):12186–12191, 2016.

[47] Qian Cong, Ivan Anishchenko, Sergey Ovchinnikov, and David Baker. Protein interaction networks revealed by proteome coevolution. Science, 365(6449):185–189, 2019.

[48] Ian R. Humphreys, Jimin Pei, Minkyung Baek, Aditya Krishnakumar, Ivan Anishchenko, Sergey Ovchinnikov, Jing Zhang, Travis J. Ness, Sudeep Banjade, Saket R. Bagde, Viktoriya G. Stancheva, Xiao-Han Li, Kaixian Liu, Zhi Zheng, Daniel J. Barrero, Upasana Roy, Jochen Kuper, Israel S. Fernández, Barnabas Szakal, Dana Branzei, Josep Rizo, Caroline Kisker, Eric C. Greene, Sue Biggins, Scott Keeney, Elizabeth A. Miller, J. Christopher Fromme, Tamara L. Hendrickson, Qian Cong, and David Baker. Computed structures of core eukaryotic protein complexes. Science, 374:1340, 2021.

[49] M. Pellegrini, E. M. Marcotte, M. J. Thompson, D. Eisenberg, and T. O. Yeates. Assigning protein functions by comparative genome analysis: protein phylogenetic profiles. Proc. Natl. Acad. Sci. USA, 96(8):4285–4288, 1999.

[50] G. Croce, T. Gueudré, M. V. Ruiz Cuevas, V. Keidel, M. Figliuzzi, H. Szurmant, and M. Weigt. A multi-scale coevolutionary approach to predict interactions between protein domains. PLOS Comput. Biol., 15(10):e1006891, 2019.

[51] Idit Bloch, Dana Sherill-Rofe, Doron Stupp, Irene Unterman, Hodaya Beer, Elad Sharon, and Yuval Tabach. Optimization of co-evolution analysis through phylogenetic profiling reveals pathway-specific signals. Bioinfor-matics, 36(14):4116–4125, 2020.

[52] D. Moi, L. Kilchoer, P. S. Aguilar, and C. Dessimoz. Scalable phylogenetic profiling using MinHash uncovers likely eukaryotic sexual reproduction genes. PLOS Comput. Biol., 16(7):e1007553, 2020.

[53] A. M. Altenhoff, C. M. Train, K. J. Gilbert, I. Mediratta, T. Mendes de Farias, D. Moi, Y. Nevers, H. S. Radoykova, V. Rossier, A. Warwick Vesztrocy, N. M. Glover, and C. Dessimoz. OMA orthology in 2021: website overhaul, conserved isoforms, ancestral gene order and more. Nucleic Acids Res., 49(D1):D373–D379, 2021.

[54] E. Dembech, M. Malatesta, C. De Rito, G. Mori, D. Cavazzini, A. Secchi, F. Morandin, and R. Percudani. Identification of hidden associations among eukaryotic genes through statistical analysis of coevolutionary transitions. Proc. Natl. Acad. Sci. USA, 120(16):e2218329120, 2023.

[55] D. Stupp, E. Sharon, I. Bloch, M. Zitnik, O. Zuk, and Y. Tabach. Co-evolution based machine-learning for predicting functional interactions between human genes. Nat. Commun., 12(1):6454, 2021.

[56] David Moi and Christophe Dessimoz. Reconstructing protein interactions across time using phylogeny-aware graph neural networks. bioRxiv, page 2022.07.21.501014, 2022.

[57] N. Konno and W. Iwasaki. Machine learning enables prediction of metabolic system evolution in bacteria. Sci. Adv., 9(2):eadc9130, 2023.

[58] Anna G Green, Hadeer Elhabashy, Kelly P Brock, Rohan Maddamsetti, Oliver Kohlbacher, and Debora S Marks. Large-scale discovery of protein interactions at residue resolution using co-evolution calculated from genomic sequences. Nature communications, 12(1):1396, 2021.

[59] A.-F. Bitbol. Inferring interaction partners from protein sequences using mutual information. PLOS Comput. Biol., 14(11):e1006401, 2018.

[60] Umberto Lupo, Damiano Sgarbossa, and Anne-Florence Bitbol. Pairing interacting protein sequences using masked language modeling. Proc. Natl. Acad. Sci. USA, 121(27):e2311887121, 2024.

[61] John Jumper, Richard Evans, Alexander Pritzel, Tim Green, Michael Figurnov, Olaf Ronneberger, Kathryn Tunyasuvunakool, Russ Bates, Augustin Žídek, Anna Potapenko, Alex Bridgland, Clemens Meyer, Simon A. A. Kohl, Andrew J. Ballard, Andrew Cowie, Bernardino Romera-Paredes, Stanislav Nikolov, Rishub Jain, Jonas Adler, Trevor Back, Stig Petersen, David Reiman, Ellen Clancy, Michal Zielinski, Martin Steinegger, Michalina Pacholska, Tamas Berghammer, Sebastian Bodenstein, David Silver, Oriol Vinyals, Andrew W. Senior, Koray Kavukcuoglu, Pushmeet Kohli, and Demis Hassabis. Highly accurate protein structure prediction with AlphaFold. Nature, 596:583–589, 2021.

[62] O. Aromolaran, D. Aromolaran, I. Isewon, and J. Oyelade. Machine learning approach to gene essentiality prediction: a review. Brief. Bioinform., 22(5):bbab128, 2021.

[63] Tristan Bepler and Bonnie Berger. Learning the protein language: Evolution, structure, and function. Cell systems, 12(6):654–669, 2021.

[64] Onkar Gujral, Mihir Bafna, Eric Alm, and Bonnie Berger. Sparse autoencoders uncover biologically inter-pretable features in protein language model representations. Proceedings of the National Academy of Sciences, 122(34):e2506316122, 2025.

[65] Carolina Rios-Martinez, Nicholas Bhattacharya, Ava P. Amini, Lorin Crawford, and Kevin K. Yang. Deep self-supervised learning for biosynthetic gene cluster detection and product classification. PLOS Computational Biology, 19(5):e1011162, 2023.

[66] David Moi and Christophe Dessimoz. Phylogenetic profiling in eukaryotes comes of age. Proc. Natl. Acad. Sci. USA, 120(19):e2305013120, 2023.

[67] Dmitry Kuznetsov, Fredrik Tegenfeldt, Mosè Manni, Mathieu Seppey, Matthew Berkeley, Evgenia V Krivent-seva, and Evgeny M Zdobnov. OrthoDB v11: annotation of orthologs in the widest sampling of organismal diversity. Nucleic Acids Res., 51(D1):D445–D451, 2022.

[68] Umberto Lupo, Damiano Sgarbossa, and Anne-Florence Bitbol. Protein language models trained on multiple sequence alignments learn phylogenetic relationships. Nat. Commun., 13:6298, 2022.

[69] Damian Szklarczyk, Rebecca Kirsch, Mikaela Koutrouli, Katerina Nastou, Farrokh Mehryary, Radja Hachilif, Annika L Gable, Tao Fang, Nadezhda T Doncheva, Sampo Pyysalo, Peer Bork, Lars J Jensen, and Christian von Mering. The STRING database in 2023: protein–protein association networks and functional enrichment analyses for any sequenced genome of interest. Nucleic Acids Res., 51(D1):D638–D646, 2022.

[70] H. M. Berman, J. Westbrook, Z. Feng, G. Gilliland, T. N. Bhat, H. Weissig, I. N. Shindyalov, and P. E. Bourne. The Protein Data Bank. Nucleic Acids Res, 28(1):235–242, 2000.

[71] Ian R. Humphreys, Jing Zhang, Minkyung Baek, Yaxi Wang, Aditya Krishnakumar, Jimin Pei, Ivan An-ishchenko, Catherine A. Tower, Blake A. Jackson, Thulasi Warrier, Deborah T. Hung, S. Brook Peterson, Joseph D. Mougous, Qian Cong, and David Baker. Protein interactions in human pathogens revealed through deep learning. Nature microbiology, 9(10):2642–2652, 2024.

[72] Jing Zhang, Ian R Humphreys, Jimin Pei, Jinuk Kim, Chulwon Choi, Rongqing Yuan, Jesse Durham, Siqi Liu, Hee-Jung Choi, Minkyung Baek, David Baker, and Qian Cong. Computing the human interactome. bioRxiv, page 2024.10.01.615885, 2024.

[73] Muhao Chen, Chelsea J T Ju, Guangyu Zhou, Xuelu Chen, Tianran Zhang, Kai-Wei Chang, Carlo Zaniolo, and Wei Wang. Multifaceted protein–protein interaction prediction based on Siamese residual RCNN. Bioinformatics, 35(14):i305–i314, 2019.

[74] Rohit Singh, Kapil Devkota, Samuel Sledzieski, Bonnie Berger, and Lenore Cowen. Topsy-Turvy: integrating a global view into sequence-based PPI prediction. Bioinformatics, 38:i264–i272, 2022. Publisher: Oxford Academic.

[75] Young Su Ko, Jonathan Parkinson, Cong Liu, and Wei Wang. TUnA: an uncertainty-aware transformer model for sequence-based protein–protein interaction prediction. Briefings in Bioinformatics, 25(5), 2024.

[76] Judith Bernett, David B Blumenthal, and Markus List. Cracking the black box of deep sequence-based protein–protein interaction prediction. Briefings in Bioinformatics, 25(2), 2024.

[77] Sanathoi Gurumayum, Puzi Jiang, Xiaowen Hao, Tulio L Campos, Neil D Young, Pasi K Korhonen, Robin B Gasser, Peer Bork, Xing-Ming Zhao, Li-jie He, and Wei-Hua Chen. OGEE v3: Online GEne Essentiality database with increased coverage of organisms and human cell lines. Nucleic Acids Research, 49(D1):D998–D1003, 2021.

[78] Clyde A. Hutchison, Ray-Yuan Chuang, Vladimir N. Noskov, Nacyra Assad-Garcia, Thomas J. Deerinck, Mark H. Ellisman, John Gill, Krishna Kannan, Bogumil J. Karas, Li Ma, James F. Pelletier, Zhi-Qing Qi, R. Alexander Richter, Elizabeth A. Strychalski, Lijie Sun, Yo Suzuki, Billyana Tsvetanova, Kim S. Wise, Hamilton O. Smith, John I. Glass, Chuck Merryman, Daniel G. Gibson, and J. Craig Venter. Design and synthesis of a minimal bacterial genome. Science, 351(6280):aad6253, 2016.

[79] Marian Breuer, Tyler M Earnest, Chuck Merryman, Kim S Wise, Lijie Sun, Michaela R Lynott, Clyde A Hutchison, Hamilton O Smith, John D Lapek, David J Gonzalez, Valérie de Crécy-Lagard, Drago Haas, Andrew D Hanson, Piyush Labhsetwar, John I Glass, and Zaida Luthey-Schulten. Essential metabolism for a minimal cell. eLife, 8:e36842, 2019.

[80] Tiago Pedreira, Christoph Elfmann, Neil Singh, and Jörg Stülke. SynWiki: Functional annotation of the first artificial organism Mycoplasma mycoides JCVI-syn3A. Protein Science, 31(1):54–62, 2022.

[81] Kai Song, Tuopong Tong, and Fang Wu. Predicting essential genes in prokaryotic genomes using a linear method: ZUPLS. Integrative Biology, 6(4):460–469, 2014.

[82] Karthik Azhagesan, Balaraman Ravindran, and Karthik Raman. Network-based features enable prediction of essential genes across diverse organisms. PLOS ONE, 13(12):e0208722, 2018.

[83] Xiao Liu, Bao-Jin Wang, Luo Xu, Hong-Ling Tang, and Guo-Qing Xu. Selection of key sequence-based features for prediction of essential genes in 31 diverse bacterial species. PLOS ONE, 12(3):e0174638, 2017.

[84] Md Abid Hasan and Stefano Lonardi. DeeplyEssential: A deep neural network for predicting essential genes in microbes. BMC Bioinformatics, 21(14):367, 2020.

[85] Adam M. Gustafson, Evan S. Snitkin, Stephen C. J. Parker, Charles DeLisi, and Simon Kasif. Towards the identification of essential genes using targeted genome sequencing and comparative analysis. BMC genomics, 7:265, 2006.

[86] Yi Yue, Chen Ye, Pei-Yun Peng, Hui-Xin Zhai, Iftikhar Ahmad, Chuan Xia, Yun-Zhi Wu, and You-Hua Zhang. A deep learning framework for identifying essential proteins based on multiple biological information. BMC Bioinformatics, 23(1):318, 2022.

[87] Soma Saha and Steffen Heber. In silico prediction of yeast deletion phenotypes. Genetics and molecular research, 5(1):224–232, 2006.

[88] Xue Zhang, Wangxin Xiao, and Xihao Hu. Predicting essential proteins by integrating orthology, gene expressions, and PPI networks. PLOS ONE, 13(4):e0195410, 2018.

[89] Yunha Hwang, Andre L Cornman, Elizabeth H Kellogg, Sergey Ovchinnikov, and Peter R Girguis. Genomic language model predicts protein co-regulation and function. Nature communications, 15(1):2880, 2024.

[90] Nishant Jha, Joshua Kravitz, Jacob West-Roberts, Cong Lu, Antonio Pedro Camargo, Simon Roux, Andre Cornman, and Yunha Hwang. Gaia: An AI-enabled genomic context–aware platform for protein sequence annotation. Science Advances, 11(25):eadv5109, 2025.

[91] Zhangzhi Peng, Benjamin Schussheim, and Pranam Chatterjee. PTM-Mamba: A PTM-aware protein language model with bidirectional gated Mamba blocks. bioRxiv, page 2024.02.28.581983, 2024.

[92] Damiano Sgarbossa, Cyril Malbranke, and Anne-Florence Bitbol. Protmamba: a homology-aware but alignment-free protein state space model. Bioinformatics, 41(6):btaf348, 2025.

[93] Jin Su, Chenchen Han, Yuyang Zhou, Junjie Shan, Xibin Zhou, and Fajie Yuan. SaProt: Protein language mod-eling with structure-aware vocabulary. In The Twelfth International Conference on Learning Representations, 2024.

[94] Michael Heinzinger, Konstantin Weissenow, Joaquin Gomez Sanchez, Adrian Henkel, Martin Steinegger, and Burkhard Rost. ProstT5: Bilingual language model for protein sequence and structure. bioRxiv, page 2023.07.23.550085, 2023.

[95] Thomas Hayes, Roshan Rao, Halil Akin, Nicholas J. Sofroniew, Deniz Oktay, Zeming Lin, Robert Verkuil, Vincent Q. Tran, Jonathan Deaton, Marius Wiggert, Rohil Badkundri, Irhum Shafkat, Jun Gong, Alexander Derry, Raul S. Molina, Neil Thomas, Yousuf A. Khan, Chetan Mishra, Carolyn Kim, Liam J. Bartie, Matthew Nemeth, Patrick D. Hsu, Tom Sercu, Salvatore Candido, and Alexander Rives. Simulating 500 million years of evolution with a language model. Science, 387(6736):850–858, 2025.

[96] Jin Su, Xibin Zhou, Xuting Zhang, and Fajie Yuan. Protrek: Navigating the protein universe through tri-modal contrastive learning. bioRxiv, page 2024.05.30.596740, 2024.

[97] Kevin Michalewicz, Mauricio Barahona, and Barbara Bravi. Integrating protein sequence embeddings with structure via graph-based deep learning for the prediction of single-residue properties. arXiv, page 2502.17294, 2025.

[98] Samuel Sledzieski, Kapil Devkota, Rohit Singh, Lenore Cowen, and Bonnie Berger. TT3D: Leveraging precom-puted protein 3D sequence models to predict protein–protein interactions. Bioinformatics, 39(11):btad663, 2023.

[99] Donald Petrey, Haiqing Zhao, Stephen J Trudeau, Diana Murray, and Barry Honig. PrePPI: A Structure In-formed Proteome-wide Database of Protein–Protein Interactions. Journal of Molecular Biology, 435(14):168052, 2023.

[100] Haiqing Zhao, Donald Petrey, Diana Murray, and Barry Honig. ZEPPI: Proteome-scale sequence-based evaluation of protein–protein interaction models. Proceedings of the National Academy of Sciences, 121(21):e2400260121, May 2024.

[101] H. Zhao, C. Velez, A. Navarene, A. Saha, J. Feldman, J. Skolnick, D. Murray, and B. Honig. Combining structural modeling and deep learning to calculate the *E. coli* protein interactome and functional networks. bioRxiv, page 2025.05.07.652715, 2025.

[102] Dapeng Xiong, Mateo Torres, Diana Murray, Le Li, Aniket C. Naravane, Robert Fragoza, Barry Honig, and Haiyuan Yu. Integrating structural homology with deep learning to achieve highly accurate protein-protein interface prediction for the human interactome. bioRxiv, page 2025.06.09.658393, 2025.

[103] Jeremy Wohlwend, Gabriele Corso, Saro Passaro, Noah Getz, Mateo Reveiz, Ken Leidal, Wojtek Swiderski, Liam Atkinson, Tally Portnoi, Itamar Chinn, Jacob Silterra, Tommi Jaakkola, and Regina Barzilay. Boltz-1: Democratizing biomolecular interaction modeling. bioRxiv, page 2024.11.19.624167, 2024.

[104] Saro Passaro, Gabriele Corso, Jeremy Wohlwend, Mateo Reveiz, Stephan Thaler, Vignesh Ram Somnath, Noah Getz, Tally Portnoi, Julien Roy, Hannes Stark, David Kwabi-Addo, Dominique Beaini, Tommi Jaakkola, and Regina Barzilay. Boltz-2: Towards accurate and efficient binding affinity prediction. bioRxiv, page 2025.06.14.659707, 2025.

[105] Josh Abramson, Jonas Adler, Jack Dunger, Richard Evans, Tim Green, Alexander Pritzel, Olaf Ronneberger, Lindsay Willmore, Andrew J. Ballard, Joshua Bambrick, Sebastian W. Bodenstein, David A. Evans, Chia-Chun Hung, Michael O’Neill, David Reiman, Kathryn Tunyasuvunakool, Zachary Wu, Akvilė Žemgulytė, Eirini Arvaniti, Charles Beattie, Ottavia Bertolli, Alex Bridgland, Alexey Cherepanov, Miles Congreve, Alexander I. Cowen-Rivers, Andrew Cowie, Michael Figurnov, Fabian B. Fuchs, Hannah Gladman, Rishub Jain, Yousuf A. Khan, Caroline M. R. Low, Kuba Perlin, Anna Potapenko, Pascal Savy, Sukhdeep Singh, Adrian Stecula, Ashok Thillaisundaram, Catherine Tong, Sergei Yakneen, Ellen D. Zhong, Michal Zielinski, Augustin Žídek, Victor Bapst, Pushmeet Kohli, Max Jaderberg, Demis Hassabis, and John M. Jumper. Accurate structure prediction of biomolecular interactions with AlphaFold 3. Nature, 630(8016):493–500, 2024.

[106] Michael Costanzo, Anastasia Baryshnikova, Jeremy Bellay, Yungil Kim, Eric D. Spear, Carolyn S. Sevier, Huiming Ding, Judice L.Y. Koh, Kiana Toufighi, Sara Mostafavi, Jeany Prinz, Robert P. St. Onge, Benjamin VanderSluis, Taras Makhnevych, Franco J. Vizeacoumar, Solmaz Alizadeh, Sondra Bahr, Renee L. Brost, Yiqun Chen, Murat Cokol, Raamesh Deshpande, Zhijian Li, Zhen-Yuan Lin, Wendy Liang, Michaela Marback, Jadine Paw, Bryan-Joseph San Luis, Ermira Shuteriqi, Amy Hin Yan Tong, Nydia van Dyk, Iain M. Wallace, Joseph A. Whitney, Matthew T. Weirauch, Guoqing Zhong, Hongwei Zhu, Walid A. Houry, Michael Brudno, Sasan Ragibizadeh, Balázs Papp, Csaba Pál, Frederick P. Roth, Guri Giaever, Corey Nislow, Olga G. Troyanskaya, Howard Bussey, Gary D. Bader, Anne-Claude Gingras, Quaid D. Morris, Philip M. Kim, Chris A. Kaiser, Chad L. Myers, Brenda J. Andrews, and Charles Boone. The Genetic Landscape of a Cell. Science, 327(5964):425–431, 2010.

[107] Michael Costanzo, Benjamin VanderSluis, Elizabeth N. Koch, Anastasia Baryshnikova, Carles Pons, Guihong Tan, Wen Wang, Matej Usaj, Julia Hanchard, Susan D. Lee, Vicent Pelechano, Erin B. Styles, Maximilian Billmann, Jolanda van Leeuwen, Nydia van Dyk, Zhen-Yuan Lin, Elena Kuzmin, Justin Nelson, Jeff S. Piotrowski, Tharan Srikumar, Sondra Bahr, Yiqun Chen, Raamesh Deshpande, Christoph F. Kurat, Sheena C. Li, Zhijian Li, Mojca Mattiazzi Usaj, Hiroki Okada, Natasha Pascoe, Bryan-Joseph San Luis, Sara Sharifpoor, Emira Shuteriqi, Scott W. Simpkins, Jamie Snider, Harsha Garadi Suresh, Yizhao Tan, Hongwei Zhu, Noel Malod-Dognin, Vuk Janjic, Natasa Przulj, Olga G. Troyanskaya, Igor Stagljar, Tian Xia, Yoshikazu Ohya, Anne-Claude Gingras, Brian Raught, Michael Boutros, Lars M. Steinmetz, Claire L. Moore, Adam P. Rosebrock, Amy A. Caudy, Chad L. Myers, Brenda Andrews, and Charles Boone. A global genetic interaction network maps a wiring diagram of cellular function. Science, 353(6306):aaf1420, 2016.

[108] Michael Costanzo, Elena Kuzmin, Jolanda van Leeuwen, Barbara Mair, Jason Moffat, Charles Boone, and Brenda Andrews. Global Genetic Networks and the Genotype-to-Phenotype Relationship. Cell, 177(1):85–100, 2019.

[109] Elena Kuzmin, Benjamin VanderSluis, Wen Wang, Guihong Tan, Raamesh Deshpande, Yiqun Chen, Matej Usaj, Attila Balint, Mojca Mattiazzi Usaj, Jolanda van Leeuwen, Elizabeth N. Koch, Carles Pons, Andrius J. Dagilis, Michael Pryszlak, Jason Zi Yang Wang, Julia Hanchard, Margot Riggi, Kaicong Xu, Hamed Heydari, Bryan-Joseph San Luis, Ermira Shuteriqi, Hongwei Zhu, Nydia Van Dyk, Sara Sharifpoor, Michael Costanzo, Robbie Loewith, Amy Caudy, Daniel Bolnick, Grant W. Brown, Brenda J. Andrews, Charles Boone, and Chad L. Myers. Systematic Analysis of Complex Genetic Interactions. Science, 360(6386):eaao1729, 2018.

[110] Enzo Kingma, Floor Dolsma, Leila Margarita Iñigo De La Cruz, and Liedewij Laan. Saturated transposon analysis in yeast as a one-step method to quantify the fitness effects of gene disruptions on a genome-wide scale. bioRxiv, page 2023.09.08.556793, 2023.

[111] Michael Costanzo, Jing Hou, Vincent Messier, Justin Nelson, Mahfuzur Rahman, Benjamin VanderSluis, Wen Wang, Carles Pons, Catherine Ross, Matej Ušaj, Bryan-Joseph San Luis, Emira Shuteriqi, Elizabeth N. Koch, Patrick Aloy, Chad L. Myers, Charles Boone, and Brenda Andrews. Environmental robustness of the global yeast genetic interaction network. Science, 372(6542):eabf8424, 2021.

[112] Michael Ashburner, Catherine A Ball, Judith A Blake, David Botstein, Heather Butler, J Michael Cherry, Allan P Davis, Kara Dolinski, Selina S Dwight, Janan T Eppig, Midori A. Harris, David P. Hill, Laurie Issel-Tarver, Andrew Kasarskis, Suzanna Lewis, John C. Matese, Joel E. Richardson, Martin Ringwald, Gerald M. Rubin, and Gavin Sherlock. Gene ontology: tool for the unification of biology. Nature genetics, 25(1):25–29, 2000.

[113] Suzi A Aleksander, James Balhoff, Seth Carbon, J Michael Cherry, Harold J Drabkin, Dustin Ebert, Marc Feuermann, Pascale Gaudet, Nomi L Harris, et al. The gene ontology knowledgebase in 2023. Genetics, 224(1):iyad031, 2023.

[114] Ashish Vaswani, Noam Shazeer, Niki Parmar, Jakob Uszkoreit, Llion Jones, Aidan N. Gomez, Łukasz Kaiser, and Illia Polosukhin. Attention is all you need. Advances in Neural Information Processing Systems, 30:5998–6008, 2017.

[115] Ahmed Elnaggar, Michael Heinzinger, Christian Dallago, Ghalia Rehawi, Yu Wang, Llion Jones, Tom Gibbs, Tamas Feher, Christoph Angerer, Martin Steinegger, Debsindhu Bhowmik, and Burkhard Rost. ProtTrans: Toward Understanding the Language of Life Through Self-Supervised Learning. IEEE Transactions on Pattern Analysis and Machine Intelligence, 44(10):7112–7127, 2022.

[116] Thomas Wolf, Lysandre Debut, Victor Sanh, Julien Chaumond, Clement Delangue, Anthony Moi, Pierric Cistac, Tim Rault, Remi Louf, Morgan Funtowicz, Joe Davison, Sam Shleifer, Patrick von Platen, Clara Ma, Yacine Jernite, Julien Plu, Canwen Xu, Teven Le Scao, Sylvain Gugger, Mariama Drame, Quentin Lhoest, and Alexander Rush. Transformers: State-of-the-art natural language processing. In Proceedings of the 2020 conference on empirical methods in natural language processing: system demonstrations, pages 38–45, 2020.

[117] Tri Dao. FlashAttention-2: Faster attention with better parallelism and work partitioning. arXiv, page 2307.08691, 2023.

[118] Diederik P Kingma and Jimmy Ba. Adam: A method for stochastic optimization. arXiv, page 1412.6980, 2014.

[119] The UniProt Consortium. UniProt: The Universal Protein Knowledgebase in 2025. Nucleic Acids Research, 53(D1):D609–D617, 2025.

[120] Eric W. Sayers, Jeffrey Beck, Evan E. Bolton, J. Rodney Brister, Jessica Chan, Ryan Connor, Michael Feldgarden, Anna M. Fine, Kathryn Funk, Jinna Hoffman, Sivakumar Kannan, Christopher Kelly, William Klimke, Sunghwan Kim, Stacy Lathrop, Aron Marchler-Bauer, Terence D. Murphy, Chris O’Sullivan, Erin Schmieder, Yuriy Skripchenko, Adam Stine, Francoise Thibaud-Nissen, Jiyao Wang, Jian Ye, Erin Zellers, Valerie A. Schneider, and Kim D. Pruitt. Database resources of the National Center for Biotechnology Information in 2025. Nucleic Acids Research, 53(D1):D20–D29, 2025.

[121] Stacia R. Engel, Suzi Aleksander, Robert S. Nash, Edith D. Wong, Shuai Weng, Stuart R. Miyasato, Gavin Sherlock, and J. Michael Cherry. Saccharomyces Genome Database: Advances in genome annotation, expanded biochemical pathways, and other key enhancements. Genetics, 229(3):iyae185, 2025.

[122] Morgan N. Price, Kelly M. Wetmore, R. Jordan Waters, Mark Callaghan, Jayashree Ray, Hualan Liu, Jennifer V. Kuehl, Ryan A. Melnyk, Jacob S. Lamson, Yumi Suh, Hans K. Carlson, Zuelma Esquivel, Harini Sadeeshkumar, Romy Chakraborty, Grant M. Zane, Benjamin E. Rubin, Judy D. Wall, Axel Visel, James Bristow, Matthew J. Blow, Adam P. Arkin, and Adam M. Deutschbauer. Mutant phenotypes for thousands of bacterial genes of unknown function. Nature, 557(7706):503–509, 2018.

[123] Martin Steinegger and Johannes Söding. Clustering huge protein sequence sets in linear time. Nature Communications, 9(1):2542, 2018.

[124] Zoe L. Watson, Fred R. Ward, Raphaël Méheust, Omer Ad, Alanna Schepartz, Jillian F. Banfield, and Jamie Hd Cate. Structure of the bacterial ribosome at 2 Å resolution. eLife, 9:e60482, 2020.

[125] Nir Kalisman, Christopher M. Adams, and Michael Levitt. Subunit order of eukaryotic TRiC/CCT chaperonin by cross-linking, mass spectrometry, and combinatorial homology modeling. Proceedings of the National Academy of Sciences, 109(8):2884–2889, 2012.

[126] F. Pazos and A. Valencia. Similarity of phylogenetic trees as indicator of protein–protein interaction. Protein Eng. Des. Sel., 14(9):609–614, 2001.

[127] D. Ochoa, D. Juan, A. Valencia, and F. Pazos. Detection of significant protein coevolution. Bioinformatics, 31(13):2166–2173, 2015.

[128] Mechthild Stoer and Frank Wagner. A simple min-cut algorithm. Journal of the ACM (JACM), 44(4):585–591, 1997.

